# Transfer RNA-derived non-coding RNAs (tncRNAs): Uncovering hidden regulators of transcriptional regulatory circuits in plants

**DOI:** 10.1101/2021.04.08.438949

**Authors:** Shafaque Zahra, Ajeet Singh, Nikita Poddar, Shailesh Kumar

**Affiliations:** National Institute of Plant Genome Research (NIPGR)

**Keywords:** tRNA, tRFs, tRNA half, non-coding RNAs, tncRNAs, plants, computational pipeline

## Abstract

The lateral emergence of distinct classes of non-coding RNAs has led to better insights into the eukaryotic regulatory networks. Among them, the existence of transfer RNA (tRNA)-derived non-coding RNAs (tncRNAs) demands their exploration in the plant kingdom. Here, we have designed a methodology to uncover the bigger picture of tncRNAome in plants. Using this, we have identified diverse tncRNAs of length 14-50 nt in ~2500 small RNA sequencing (sRNA-seq) samples of six major angiosperms, and further studied their various features including length, codon-usage, cleavage pattern, and modified tRNA nucleosides. Codon-dependent generation of tncRNAs indicates that the process is highly specific rather than being mere random tRNA degradation. Analysis for nucleotide composition of tncRNA cleavage positions indicates that they are generated through precise endoribonucleolytic machinery. Certain tRNA nucleoside modifications on tncRNAs were found to be conserved across the plants, and hence may influence tRNA cleavage, as well as tncRNA functions. Pathway enrichment analysis revealed that common tncRNA targets were majorly involved in the metabolic and developmental processes of plants. Also, many tRFs were found to be associated with transposable elements. Distinct tissue-specific tncRNA clusters indicate their role in plant development under normal physiological conditions. Furthermore, the identification of a significant number of differentially expressed tncRNAs under several abiotic and biotic stresses highlights their probable role as gene expression modulators during various stress conditions. Thus, this study will be beneficial to investigate the emerging role of tncRNAs as prospective biomarkers in plant development and stress.

**Highlights:**

- Computational pipeline for accurate identification of genuine transfer RNA-derived non-coding RNAs (tncRNAs) using small RNA sequencing (sRNA-seq) datasets.
- Six major tncRNA classes of length ranging from 14 to 50 nt were identified in ~2,500 sRNA-seq datasets in six different angiosperms.
- tRNA nucleoside modifications may affect tncRNA cleavage in plants.
- Conserved tncRNAs target transcripts are involved in plant growth, development, and metabolism.
- tncRNAs are expressed in a tissue-dependent, and stress-specific manner in plants.

## 1. Introduction

The profusion of non-coding RNAs (ncRNAs) with diverse regulatory actions discovered over the last decade has proved to be a crucial breakthrough in the field of molecular biology. They are powerful regulators of gene expression at the epigenetic, transcriptional, and post-transcriptional levels in the living system [1]. In this diverse pool of smaller ncRNAs, miRNAs and siRNAs are the most extensively surveyed molecules [2]. They alter gene expression by binding to their target mRNAs [3]. The advancement of high-throughput sequencing technology has tremendously resolved the surveillance of the other classes of small untranslated RNAs beyond miRNAs and siRNAs [4]. Amongst them, transfer RNA (tRNA)-derived non-coding RNAs (tncRNAs) have been reported in all three domains of life [5]. This distinct group of regulatory RNAs is derived from the endonucleolytic cleavage of precursor or mature tRNAs [6]. tncRNAs include well-known shorter tRNA-derived RNA fragments up to 30 nt lengths popularly termed as tRFs [7] (or tDRs [8]) and longer tRNA halves [9] (tRHs; also recognized as tRNA-derived stress-induced fragments [10] or tiRNAs) that are usually longer than 30 nt. Precise tRNA cleavage leading to their production is governed by tRNA type, cell type, developmental stage, stress, and tissue [11]–[13]. It has been revealed that these novel molecules can originate from both nuclear as well as organellar tRNAs [14], [15]. This class of small RNAs in the range of 12-50 nucleotides (nt) has been identified by various research groups and has been particularly well studied in human cancers [16]–[18]. Several reports have demonstrated the tRFs as global regulators of gene expression because of their association with Argonaute (AGO) proteins [19]–[21]. They are frequently dysregulated in multiple types of human cancer and numerous other ailments like viral infections [22], neurodegenerative [23], and metabolic disorders [24], [25]. These fragments are speculated to govern diverse cellular processes such as cell proliferation [6], translation initiation [26], [27], stress granule assembly [23], ribosome biogenesis [28], and apoptosis [29].

In plants, the earliest report by Thompson et al in 2008 showed that the processed tRNA halves were generated in *Arabidopsis* subjected to oxidative stress [30]. In one work, phloem-specific tRNA fragments were shown to interfere with protein synthesis in pumpkin [31] (*Cucurbita maxima*). In *Arabidopsis*, a group of 19 nt small RNAs derived from tRNAs in addition to longer tRNA halves corresponding to the 5’ end of tRNA, and expressed at a high level in phosphate-starved roots [32]. For the first time in plants, Loss Morais and his group in 2013 showed the association of 19 nt tRFs with Argonaute by utilizing deep sequencing libraries of immunoprecipitated AGO proteins under stress conditions [33]. Alves et al. extended the research and highlighted the differential accumulation of specific tRFs under abiotic stresses in *Arabidopsis* and rice [34]. In addition to abiotic stresses, some studies have shown links to tncRNAs generation in response to biotic stresses e.g. Apple stem grooving virus (ASGV) infected domesticated apple trees [35] and black pepper infected with *Phytophthora capsici* [36].

Among the eukaryotic tncRNA population, shorter tRFs range from ~ 14-30 nt while the longer tRHs are ~30-40 nt in size [11], [37]. Depending on the position of cleavage on mature tRNA, there are two types of tRFs: 5’ end derived tRF-5, and 3’ end derived tRF-3 generated upon cleavage in the D and T region respectively. Among the tRNA halves, 5’ or 3’ tRHs are produced by cleavage in the anticodon region containing 5’ or 3’ portions of mature tRNA [38]. The precursor tRNA produces 5’U-tRFs/leader tRF, and tRF-1/tsRNA/3’U-tRFs from the 5’ leader and 3’ trailer portion respectively [39]–[41]. By now, it has been established that tncRNAs can be generated in a DICER-dependent or independent manner in plants [42], [43]. Besides Dicer-like proteins (DCLs), specific endoribonucleases, RNS belonging to the RNase T2 family are the major contributors to the biogenesis of tncRNAs originating from mature tRNAs [34]. Both shorter and longer tncRNAs are produced by RNS1 in *Arabidopsis* and are evolutionarily conserved across the plant kingdom [44]. The tncRNA generation from precursor tRNAs in plants has been understudied. The nomenclature of tncRNA subcategories is based on the cleavage site of their progenitor tRNAs although a homogenized nomenclature is still lacking.

Identification of tncRNAs in small RNA-seq datasets is a very challenging and error-prone process. A major reason is the extensively modified tRNA nucleosides affect the reverse transcription process during library preparation leading to truncation or misincorporation of nucleotides [45]. This increases the probability of mismatch, thereby hampering the sequencing results. During the mapping of sequenced reads, allowing just a single mismatch to overcome sequencing errors, may also lead to base misidentification and hence causes the inflation of the false negatives. This is because of high sequence similarities among 20 major tRNA isotypes covering the whole tRNA family of isoacceptors (tRNAs with different anticodons but charged with the same amino acid), and isodecoders (tRNAs carrying the same anticodon with variations in the body sequence). Also, mapping reads to both genome and tRNA is necessary for accurate tncRNA prediction as mapping to tRNA set alone may increase the chance of ambiguous fragments being identified as tncRNAs. To overcome these discrepancies, different mapping strategies for tRNA-derived reads have been suggested by researchers [46]–[48]. Some tools are available for tncRNA detection e.g. tDRmapper [49], MINTmap [50], and tRF2Cancer [51], but they are trained and tested on human datasets, and thereby suit for human data only. As compared to humans, tncRNAs have been less explored *in planta*. Although, plant exclusive databases e.g. tRex [52] and PtRFdb [53] harboring information related to the transfer RNA-derived fragments (tRFs), have also been developed in the recent past yet a convenient methodology for the accurate identification and exploration of tncRNAs is still lacking in plants.

In this study, we have developed a computational methodology for the accurate identification of tncRNAs in small RNA sequencing datasets and identified different kinds of tncRNAs in six different Angiosperms viz. *Arabidopsis thaliana* (model plant), *Solanum lycopersicum* (tomato), *Cicer arietinum* (chickpea), *Medicago truncatula* (model legume), *Oryza sativa* (rice), and *Zea mays* (maize) by exploiting publicly available small RNA sequencing datasets. Using our in-house built pipeline, we have identified, cataloged, and quantified tncRNAs into different subtypes based on their origin and cleavage position viz. tRF-5, tRF-3, tRF-1, leader tRF, 5’ and 3’ tRNA halves (tRHs). As there is no single standardized nomenclature for tncRNA subtypes yet, we have followed the classical and the most frequently used terms by researchers for tncRNA subtypes. tRNA modifications play an integral role in tRNA maturation, structural integrity, and stabilization in all domains of life [54], [55]. These modified nucleosides are also involved in a diverse range of biological activities such as gene expression regulation, growth, development, and response to stresses [56], [57]. Recently, few studies have indicated that some specific tRNA modifications may drive programmed tRNA cleavage thus leading to the genesis of tncRNAs [58], [59]. Thus, in addition to the tncRNA identification, we have also incorporated information related to the type of modified nucleosides detected, and their respective position on the individual identified tncRNAs in our analyzed samples. We also scrutinized tissue-specific expression of various tncRNA classes from distinct tissues in *Arabidopsis*. It suggests the role of tissue-specific tncRNA production under normal physiological conditions. Furthermore, we have also cataloged the significant differentially up- and down-regulated tncRNAs under various abiotic, and biotic stresses from ~1000 libraries from two dicots (*Arabidopsis* and tomato) and two monocots (rice and maize) plants. By using our methodology, this study has highlighted a major class of ncRNAs derived from tRNA in plants. In addition to this, this study also provides a future perspective for better understanding the role of tRNA modifications in tRNA biology. Differences in tncRNA expression in various conditions suggest their multiple role in plant physiology in relevance to both normal and stress conditions.

## 2. Results

### 2.1. *In silico* tncRNA identification across six plants

We have identified tRNA-derived non-coding RNAs in six angiosperms by utilizing 2,660 small RNA sequencing samples available at Sequence Read Archive (SRA), from the National Center for Biotechnology Information (NCBI). We downloaded the available datasets for *A. thaliana* (1832), *S. lycopersicum* (175), *C. arietinum* (25), *M. truncatula* (139), *O. sativa* (258), and *Z. mays* (231). After processing and quality control, filtered reads were provided as input to the in-house designed pipeline for precise tncRNA identification (Fig. 1A). Due to the huge number of tRNA gene copies predicted over the entire genome by tRNAscanSE [60], we have filtered our tRNA gene pool by eliminating pseudogenes and keeping genuine tRNAs based on score. The score is an important indicator of the structural propensity of the predicted tRNA [60], [61]. Thus, only high-quality tRNAs with a score equal to or above 50 were selected for mapping. To avoid ambiguous reads coming from non-tRNA regions, we created an artificial genome, by masking the genuine tRNA gene region with 50 nt upstream and downstream in the genome and adding them as artificial chromosomes. We classified only those reads as tncRNAs that mapped exclusively to the artificial chromosomes. The identified tncRNAs were categorized into tRF-5, tRF-3, tRF-1, leader tRF, 5’tRH, and 3’tRH (Fig. 1B). Apart from them, we have also segregated the reads mapping to the internal regions of mature tRNA and collectively grouped them under other tRFs (Fig. 1B). The various information related to each tncRNA i.e. tncRNA type, parent tRNA locus information, position (start-end), sequence, length, read count, RPM, nature of modified nucleoside(s), and its position has been provided in the output file for each SRA sample. Additionally, the alignment of each tncRNA on its progenitor tRNA can be visualized in an alignment file. The count of different tncRNA subclasses for each sample is also provided in a log file.

**Figure 1:**
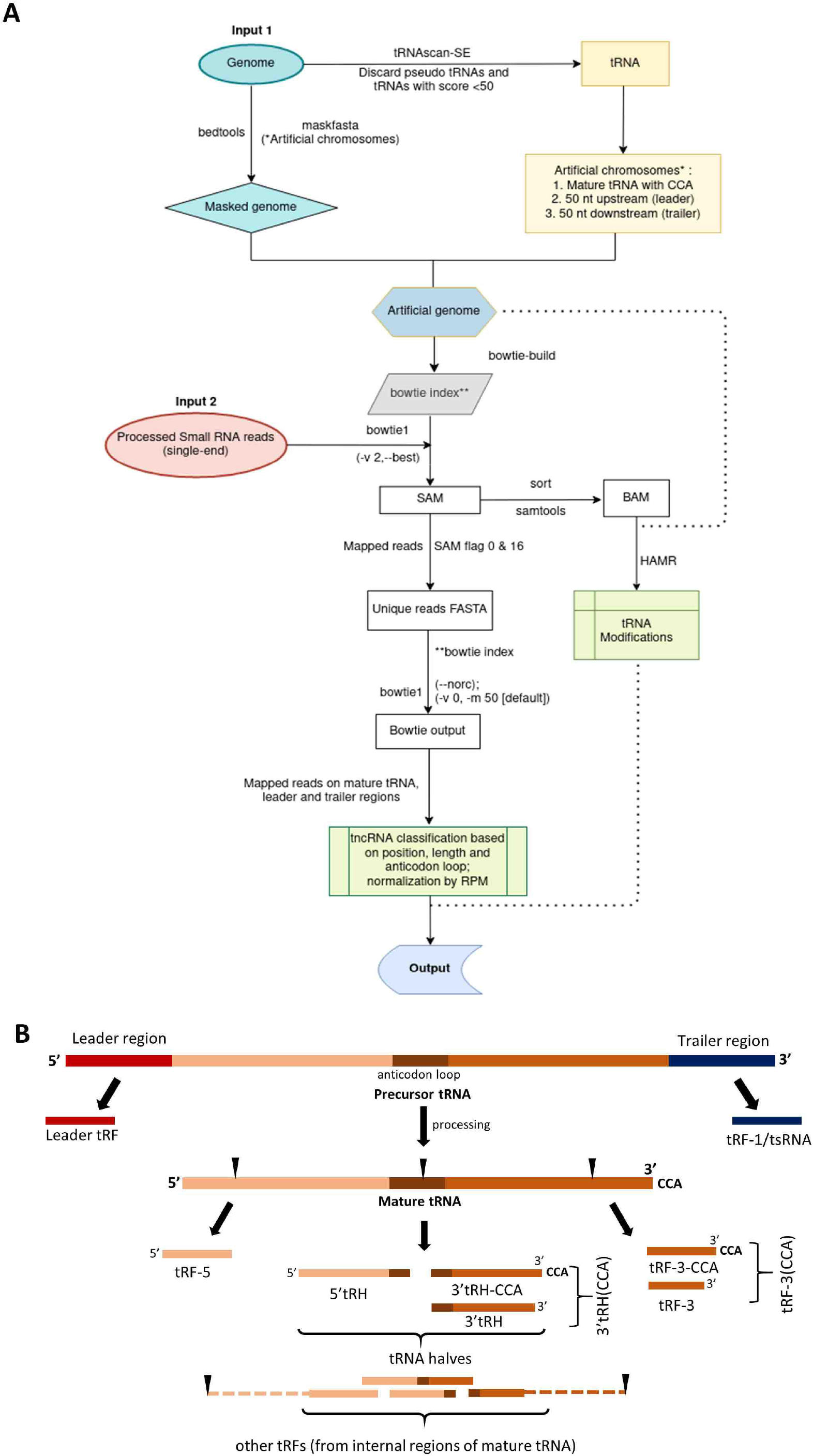
Identification of transfer-RNA-derived non-coding RNAs (tncRNAs). **A** Workflow for the identification of various classes of tncRNAs using reference genome, and processed small RNA-seq data based on tRNA origin, and site of cleavage. **B** Biogenesis of distinct tncRNA subtypes generated from the excision of precursor, and mature tRNA. pre-tRNA gives rise to leader tRF and tRF-1/tsRNA, whereas mature tRNA generates tRF-5, tRF-3(CCA), 5’tRH, 3’tRH(CCA), and other tRFs (derived from internal portions excluding extreme ends of mature tRNA denoted by dashed lines).

For the analysis, we considered only those samples that generated a minimum of 100 tncRNAs, and our analyses comprise tncRNAs from 2,448 RNA-seq samples viz. *A. thaliana* (1676), *S. lycopersicum* (160), *C. arietinum* (21), *M. truncatula* (127), *O. sativa* (243), and *Z. mays* (221). The total, as well as unique tncRNAs (the same tncRNA sequences were grouped and termed as unique), counts for each plant have been summarised in Table 1. The relative percentage of sRNA samples, tncRNA abundance (total and unique), tncRNA classes (total and unique) for each plant are shown in Fig. 2A. The samples from *Arabidopsis* outnumbered other plants and nearly 70% of all identified total tncRNAs belonged to the model plant. Chickpea samples were the least in number, hence the tncRNAs from this plant comprised only 0.01% of the total identified tncRNAs. We also observed that the relative abundance of unique sequences is fairly comparable to *Arabidopsis* in other plants (the removal of duplicate sequences coming from several samples reduces the bias of sample number). Although ~70% of samples were from *Arabidopsis*, unique tncRNAs from *Arabidopsis* contributed to 35% of the total tncRNAs. The unique sequence count per tncRNA category for individual samples from each plant has been provided in Supplementary sheet 1 (Sheets 1.1-1.7). The generation of tncRNAs showed a weak positive correlation (Pearson correlation coefficient ranged from 0.21-0.56) with the abundance of tRNA gene copies (Supplementary Figure 1). This supports the fact that the generation of tncRNAs is a non-random event and the number of tncRNAs generated is not directly proportional to tRNA frequency.

**Figure 2:**
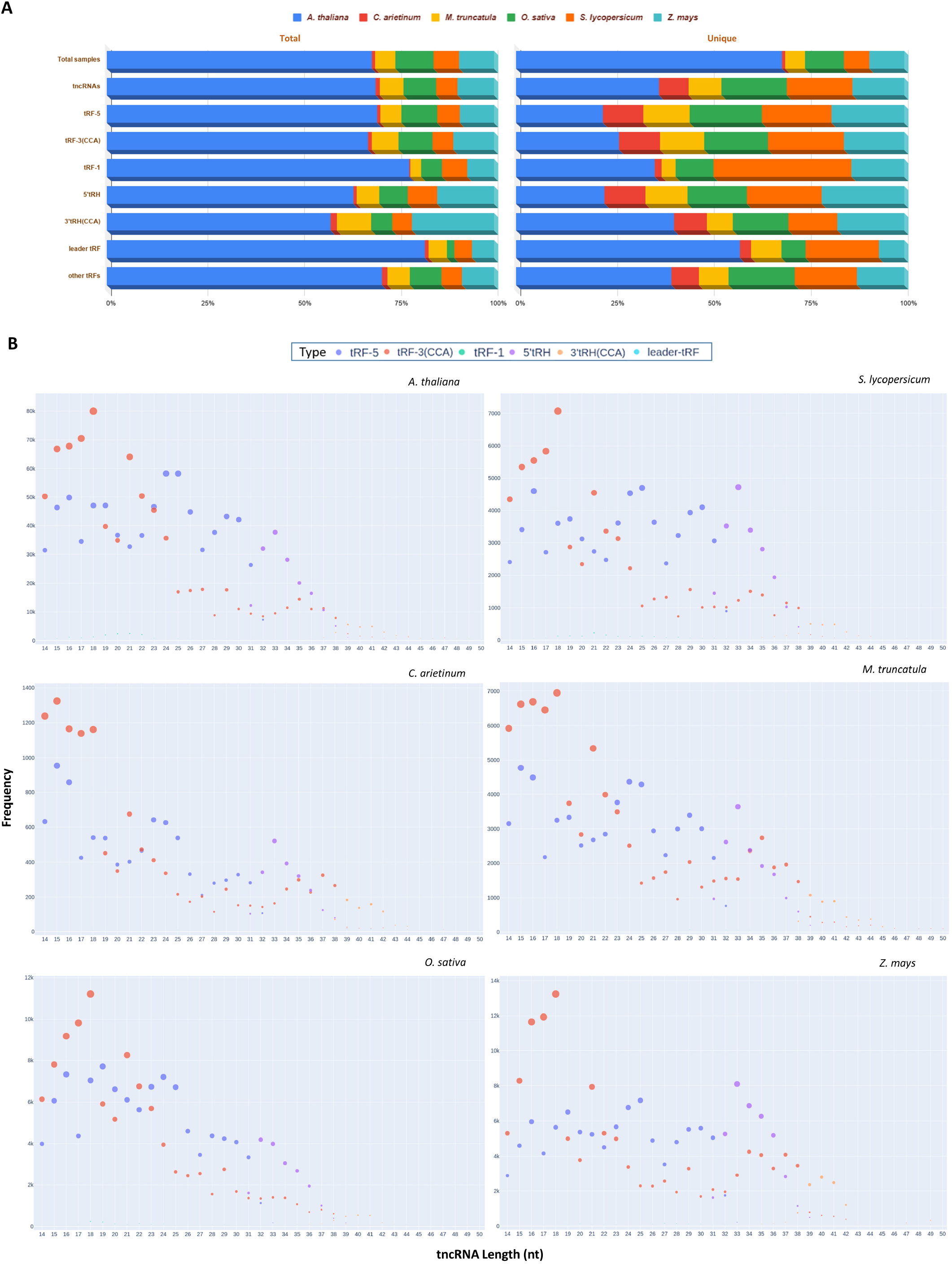
Abundance of the identified transfer-RNA-derived non-coding RNAs (tncRNAs) in different plants. **A** Relative distribution of samples, total and unique tncRNA, tRF-5, tRF-3(CCA), tRF-1, 5’tRH, 3’tRH(CCA), leader, and other tRFs analyzed in this study. **B** Size distribution of major tncRNA categories viz. tRF-5, tRF-3(CCA), tRF-1, 5’tRH, 3’tRH(CCA), and other tRFs in individual plants.

**Table 1:**
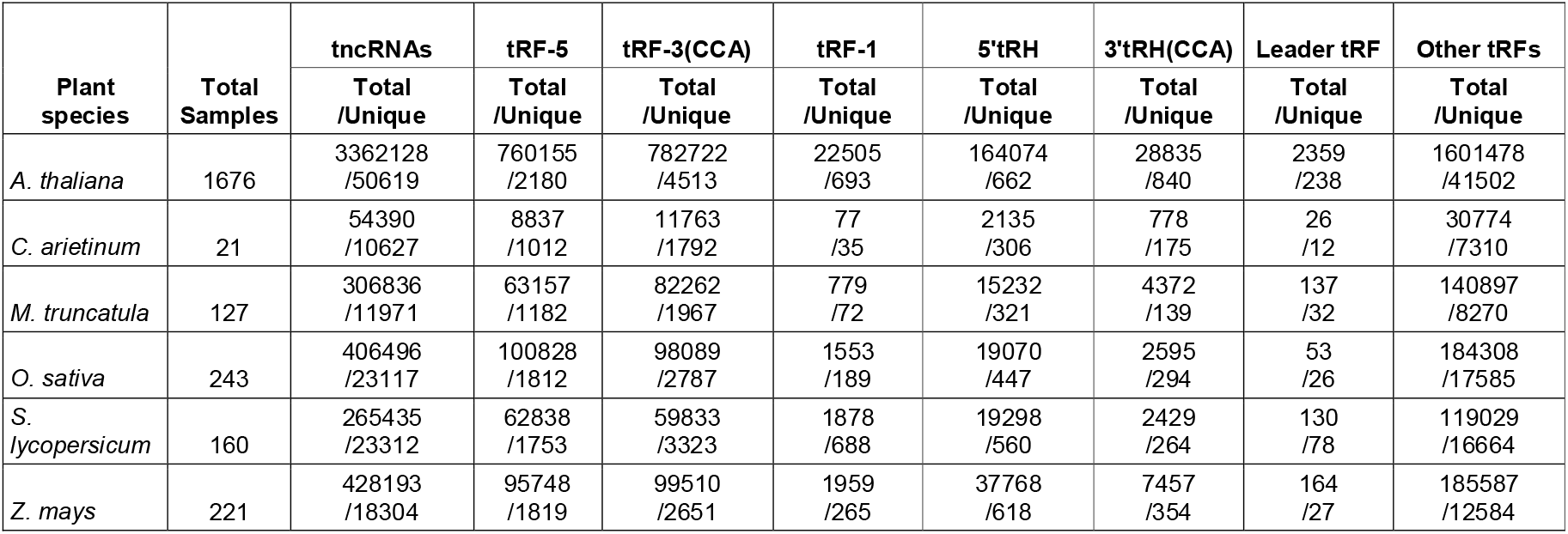
Summary of identified transfer-RNA-derived non-coding RNAs (tncRNAs) abundance in six plants.

### 2.2. tncRNA classification according to origin and site of cleavage

The tncRNA classes ranging from 14-50 nt were identified, and their length-wise abundance in all six plants (Fig. 2B). The tncRNAs belonging to the tRF-5 series ranged from ~14-33 nt, and showed widespread distribution of varying lengths in all six plants. These fragments containing the 5’ end of mature tRNA are mostly generated by cleavage in the D region. Some specifically sized fragments showed very high read counts e.g. in *Arabidopsis*, 24-25 nt long tRF-5s were most recurrent in the total tRF-5 population. The tncRNAs belonging to the tRF-3(CCA) series derived from the 3’ end of mature tRNA by cleavage in the T region (with or without CCA at 3’ end) were observed to be falling in the broad range of 14-50 nt. However, the bulk of the identified tncRNAs belonging to the tRF-3 series fall in a smaller length range up to 21 nt. The high frequency of specific-sized tncRNAs indicates cleavage preference by the endoribonuclease machinery at a certain position in the loop/stem region. Also, more than 92% of the total identified tRF-3 sequences contained terminal end CCA in all plants. The fragments corresponding to the longer halves were generated by cleavage in the anticodon region of mature tRNA. The 5’ tRHs containing the terminal 5’ region ranged from 31-40 nt (Fig. 2B). The 3’tRH(CCA) containing the terminal 3’ portion was longer than 5’ tRHs and ranged from 35-50 nt. Greater than 80% of them contained CCA at the 3’ end. The high abundance of CCA ending fragments indicates that the terminal CCA might also play a role in the structural and functional integrity of fragments generated from the 3’ end i.e. tRF-3 series and 3’ tRNA halves.

As compared to tncRNAs derived from mature tRNAs, the tncRNAs generated from the trailer and leader sequences occupied a very tiny percentage of the total tncRNA pool. The highly recurrent tRF-1 fragments fall in the range of 18-22 nt in *Arabidopsis*, tomato, rice, and maize. Lowly expressed longer tRF-1 fragments of varying size (25-50 nt) were also detected in different plants. Fragments derived from the leader region were of varying length ranging from 15-50 nt. In *Arabidopsis*, ~20 nt and longer (e.g. 48-50 nt length) fragments were recurrent. Among various tncRNAs identified, greater than 40% of the reads belonged to tRF-5 and tRF-3 series in individual plants (Supplementary Figure 2). We also observed a remarkable fraction of fragments generated from internal portions of mature tRNA. These fragments constitute nearly 45% of the total tncRNA sequences identified in each plant (Supplementary Figure 2). Longer tRHs occupy less than 10% of the tncRNAs. Compared to 3’tRHs, the 5’ tRHs were more abundant. The remaining fraction (<1%) comprised of tncRNAs generated from pre-tRNAs.

Our observations confirm the existence of a heterogeneous pool of tncRNAs in plants. In the tncRNA pool, large numbers of identical tncRNAs differing only by a few base pairs were also observed in high numbers. It is still unclear how size drives the function of tncRNAs. The tncRNA length specificity may have a structure-function relationship. The length diversity of tncRNAs indicates their prospective roles besides canonical miRNA-like gene expression regulation. The high abundance of tncRNA of a specific length or falling within a narrow length range, e.g. 16 nt tRF-5 and tRF-3 sequences need to be carefully examined as smaller fragments are usually discarded or considered to be randomly degraded fragments. The highly abundant and diverse reads belonging to tRF-5, tRF-3, as well as those derived from the internal region of tRNAs, suggest that they may have varied roles in plants whereas longer tRNA halves may be only generated during specific conditions. Although observed in low frequency than other tncRNA categories, we have observed tRF-1 expression in various samples. Till now in plants, both tRF-1 and leader tncRNAs remain unidentified because of their very low abundance and demands to be extensively explored.

### 2.3. Cleavage of tncRNAs is highly specific and conserved amongst plants

Nucleotide composition and the ratio of the constituent monomer units is an important characteristic feature for studying nucleic acids. For studying the tncRNA cleavage pattern, we looked into the nucleotide composition of the tncRNA::tRNA junction for highly recurring tRF-5, tRF-3(CCA), 5’tRH, and 3’tRH(CCA) sequences (Fig. 3). A common conserved motif was observed in tRF-5::tRNA interface among all plants i.e. AG::TGG (first row in Fig. 3) in tRF-5s ranging between 15-19 nt. Also, the last nucleotide in tRF-5 was observed to be A/G rich (first three rows from top in Fig. 3). In tRNA::tRF-3(CCA) junction, CGA(N)::(N)TC pattern was commonly observed across all species (fourth to sixth rows in Fig. 3). Similarly, in the majority of the 5’ tRNA halves, the breakpoint region in the middle (fifth and sixth position) is either T/C (seventh to ninth rows in Fig. 3) whereas the first nucleotide in 39 nt 3’ tRH(CCA) is A rich in all species (sixth nucleotide in the tenth rows in Fig. 3). Also, a common conserved pattern, CTTGTAAAC was observed in a majority of 3’tRHs of 40-41nt length (eleventh and twelfth rows in Fig. 3). Despite the variability in the anticodon region due to anticodon triplets in tRNA halves coming from various isoacceptors, we have seen common motifs among different plants indicating that specific isoacceptors are mostly preferred by the cleaving enzymes. Based on observed motifs and specific nucleotide composition of tncRNAs, it is worth speculating that the enzymatic cleavage might first recognize or prefer a particular motif located on parental tRNA to cleave precise fragments for at least a reasonable number of genuine tncRNAs. Conservation of cleavage consensus sites in different plants supports that they are the product of non-random ribonucleolytic tRNA cleavage. Not only for their generation, but they may also aid in their stability and interaction with their cognate binding partners. It has been reported that G at 18^th^ and 19^th^ position in tRF-5 plays a crucial role in the inhibition of protein translation in both human and *Arabidopsis* [62], [63]. Similarly, motif-dependent translation inhibition by 5’tRH was also observed [62]. It can be speculated that at least some of the conserved residues may also play a synergistic role in governing tncRNA functions.

**Figure 3:**
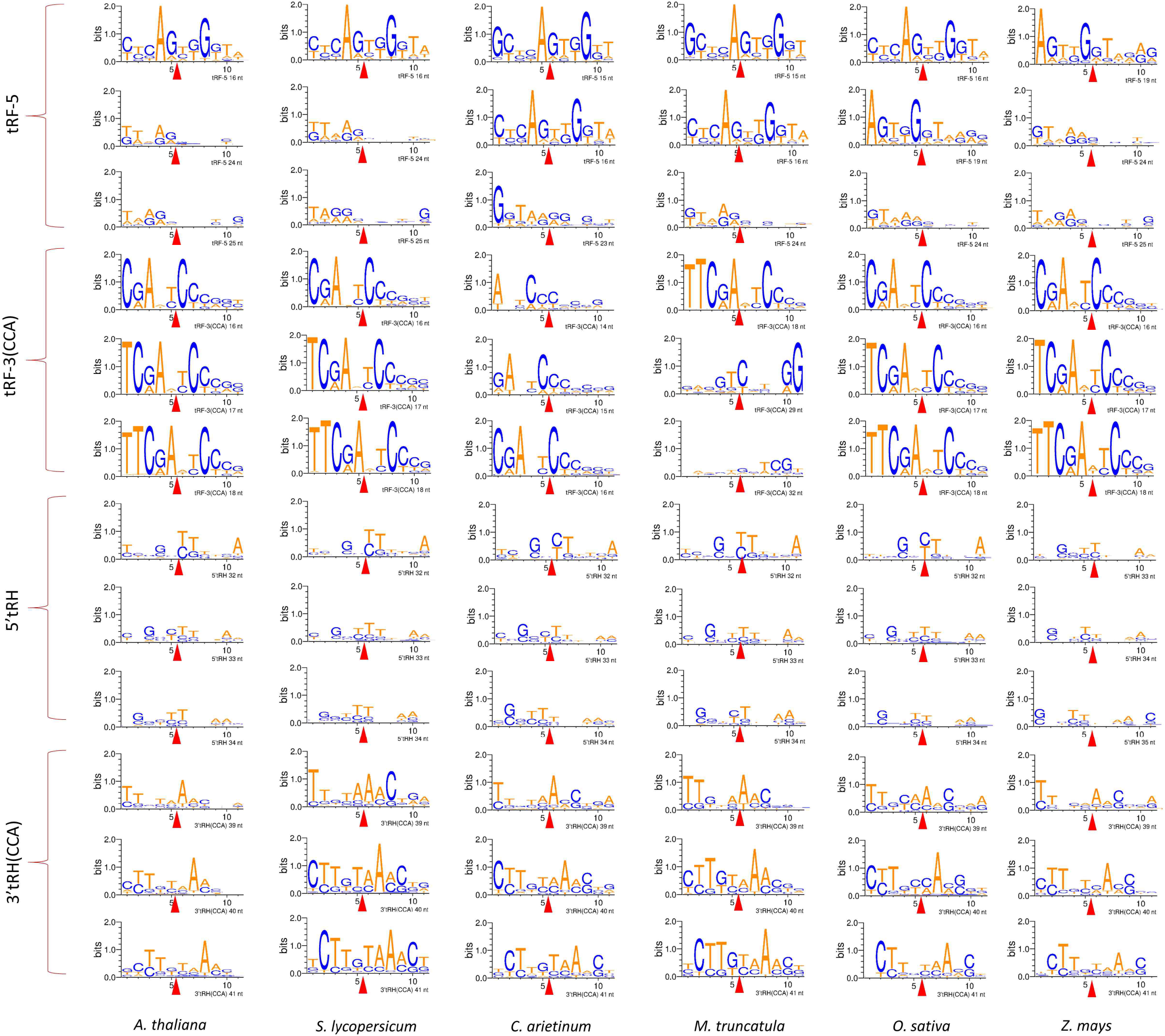
Web logo representation depicting conserved cleavage sites for four tncRNA subclasses viz. tRF-5, tRF-3(CCA), 5’tRH, and 3’tRH(CCA). Web Logos were made using top-three recurring tncRNAs by taking 5 nt stretches from progenitor tRNA and respective tncRNA fragments viz. tRF-5::tRNA; 5’tRH::tRNA; tRNA::tRF-3(CCA);tRNA::3’tRH(CCA). To study the breakpoint and its flanking nucleotide composition, for tRF-5 and 5’tRH classes, the first five nucleotides (from left to right) represent the last five nucleotides of tRF-5/5’tRH and the remaining stretch of nucleotides is the tRNA (sixth to the eleventh nucleotide). For tRF-3(CCA) and 3’tRH(CCA) classes, the first five nucleotides (from left to right) represent progenitor tRNA sequence, and the remaining stretch represents the first sixth nucleotide of tRF-3(CCA)/3’tRH(CCA) (sixth to the eleventh nucleotide). The red arrow represents the breaking point of each tncRNA fragment from its parental tRNA. All the sequences shown are oriented in the 5’ to 3’ direction. G|T; T|C; A|A breakpoint was commonly found in all types of tncRNAs.

### 2.4. Specific anticodons, not the whole tRNA repertoire are responsible for tncRNA generation

Out of the 64 triplet codons, 61 are sense codons that code for 20 amino acids. We observed that not all 61 codons generated tncRNAs. Out of 61 codons, *Arabidopsis* utilized 49, chickpea:41, *Medicago:44*, while 48 sets of tRNA isoacceptors were utilized by tomato, rice, and maize for the generation of different tncRNAs (Fig. 4). Different isoacceptors contributed to the production of different tncRNA types and varied from one plant to another. Although coding for the same amino acids, there was marked variation among the isoacceptors for contributing to the tncRNA generation e.g. in *Arabidopsis*, among five tRNA-Arg isoacceptors viz. ACG, CCG, CCT, TCG, and TCT, tRNA-ArgACG generated the most number of tRF-5s. Also, many anticodons may give rise to more than one class of tncRNAs e.g. GluCTC, ProCGG contributed for tRF-5 and tRF-3 production in *Arabidopsis*. Similarly, GluCTC, GlyTCC, LeuTAA generated both 5’tRHs and 3’tRHs in *Arabidopsis*. Also, cleavage in the anticodon loop can generate both types of tRNA halves. Whether a single cleavage in the tRNA molecule can generate functional 5’ and 3’ tRH simultaneously is unclear. There must be implications pertaining to tRNA isoacceptors and the binding of cognate cleaving machinery for driving the tncRNA generation which is currently unexplored. Moreover, there can be many reasons for codon-dependent generation of tncRNAs, such as different conditions of plant growth, developmental stages, abiotic or biotic stress, etc., and also can vary from species to species.

**Figure 4:**
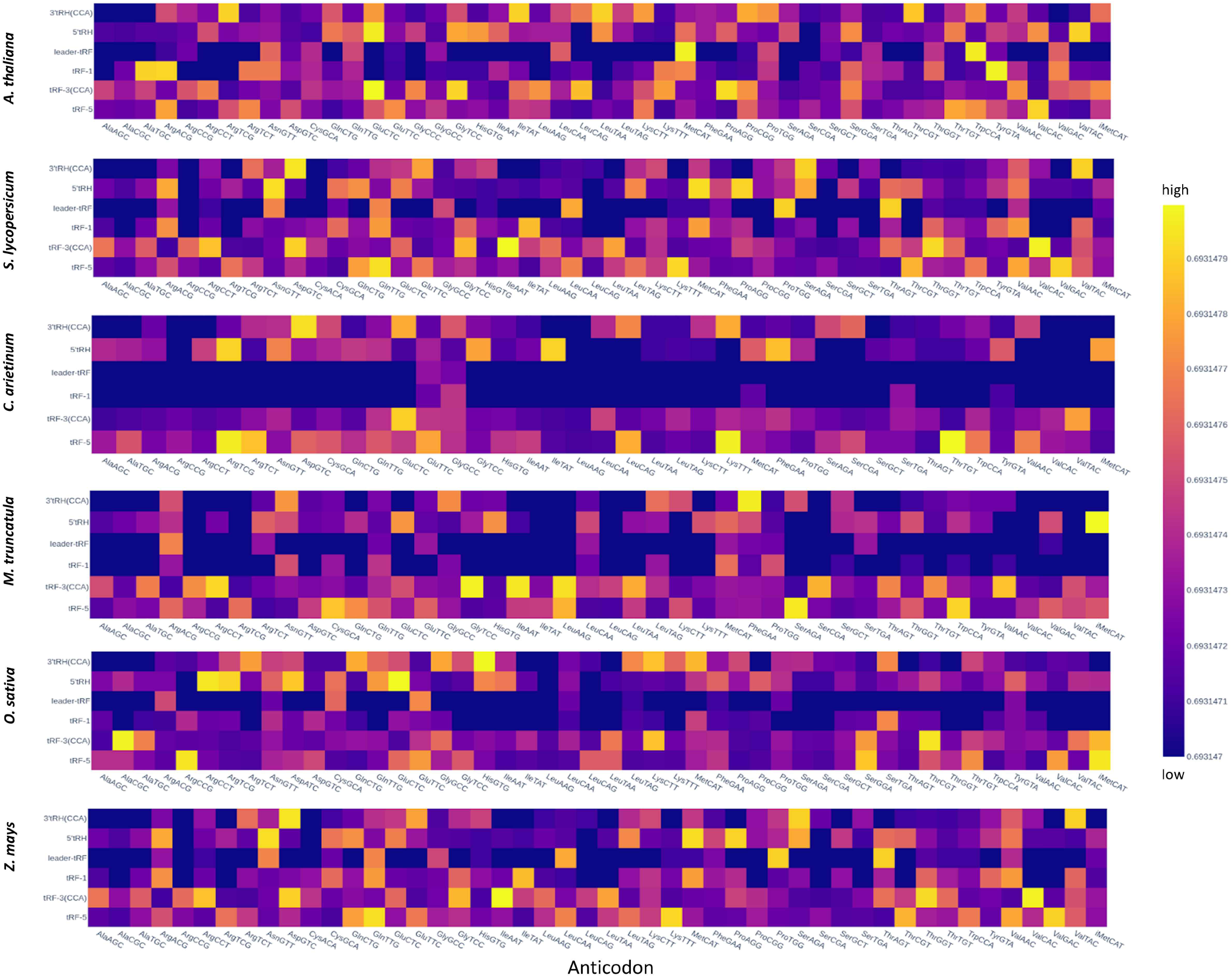
Heatmap showing the abundance of tncRNA subclasses viz. tRF-5, tRF-3(CCA), tRF-1, 5’tRH, 3’tRH(CCA), and leader tRF sourcing from each tRNA isoacceptor (specified using their respective anticodon) in six plants; The frequencies are given in read count converted to log10 values.

### 2.5. Modification may play a major role in regulating the generation, stability, and function of tncRNAs

As tRNAs are heavily modified molecules with over 100 known modifications, we scrutinized the modifications present in our identified tncRNAs. For this, we utilized HAMR[64] which is a high utility tool for the identification of modified nucleosides in high throughput sequencing datasets. Various modifications were detected by HAMR on tncRNAs in different plants viz. 3-methylcytosine (m^3^C), pseudouridine (Y/Ψ), dihydrouridine (D), N^*2*^-methylguanosine (m^2^G), N^*2*^-dimethylguanosine (m^2^_2_G), 1-methyl guanosine (m^1^G), 1-methyladenosine (m^1^A), 1-methylinosine (m^1^I), 2-methylthio-N^6^-isopentenyladenosine (ms^2^i^6^A), N^6^-isopentenyl adenosine (i^6^A), Threonylcarbamoyladenosine (t^6^A) in various isoacceptors generating tncRNAs. The visualization of various modifications on tncRNAs with respect to their position on the respective consensus sequence of mature tRNA in 49 isoacceptors in *Arabidopsis* has been shown in Supplementary Figure 3. For other plants individually, it has been provided in Supplementary Figure 4-8. We observed that the majority of modifications were detected from 20^th^ to 40^th^ position and 50^th^ to 60^th^ position i.e. containing the cleavage sites for tRF-5, tRHs, and tRF-3, indicating a high probability of tncRNA generation to be regulated by tRNA modifications. The recurring tRNA modifications in tncRNAs from *Arabidopsis* samples have been highlighted in Supplementary Figure 9.

It was observed that some modifications were specific for certain anticodons and were found in abundance at specific positions. E.g. in the D region, Guanosine (G) was modified to m^1^G on GlyGCC, and iMetCAT (Supplementary Figure 9). Also pseudouridine (Y) modification on GlyGCC tRNAs was highly abundant. In the anticodon loop region, Y and D were abundant in tncRNAs originating from GluTTC tRNAs. In the T region, m^1^A|m^1^I|ms^2^i^6^A were observed in a large number of tRNAs. We also looked for conserved modifications on different isoacceptors found common in all six plants selected for this study (Fig. 5). Adenosine (A) replaced with m^1^A|m^1^I|ms^2^i^6^A modifications were found in 20 isoacceptors viz. tRNA ArgACG|CCG|TCG|TCT (58^th^ position), AspGTC (57^th^ position), GlnTTG (57^th^ position), GlyGCC (57^th^ position), iMetCAT (57th position), LeuCAA (69^th^ position), LeuCAG (66^th^ position), LeuTAG (65^th^ position), LysCTT (58^th^ position), LysTTT (57^th^ position), SerAGA|CGA|TGA (67^th^ position), ThrTGT (57^th^ position), TyrGTA (58^th^ position), and ValAAC|CAC (59^th^ position). G is modified to m^1^G on the T loop at 58^th^ position in TyrGTA and 59^th^ position in ValAAC|CAC. These modifications were prominent in the region of the T loop which is the site of cleavage for tRF-3 generation. Also, on the 9^th^ nucleotide, G was replaced with m^1^G in ArgACG, GlyGCC, and iMetCAT in all plants. Various tncRNAs have been associated with abiotic stresses. E.g. Am nucleoside (2’-O-methyladenosine) is induced by salt stress in a variety of land plants including *Arabidopsis, Brachypodium*, poplar, and rice[65]. 2’-O-methylguanosine (Gm), 5-methyluridine (m^5^U), and 5-methylcytidine (m^5^C) are also tightly linked to plant development [57]. The findings indicate the possibility of a very delicate cross-talk among the genetic machinery involved in tncRNA biogenesis, functioning, as well as tRNA nucleoside modification in response to plant growth, development, as well as environmental changes.

**Figure 5:**
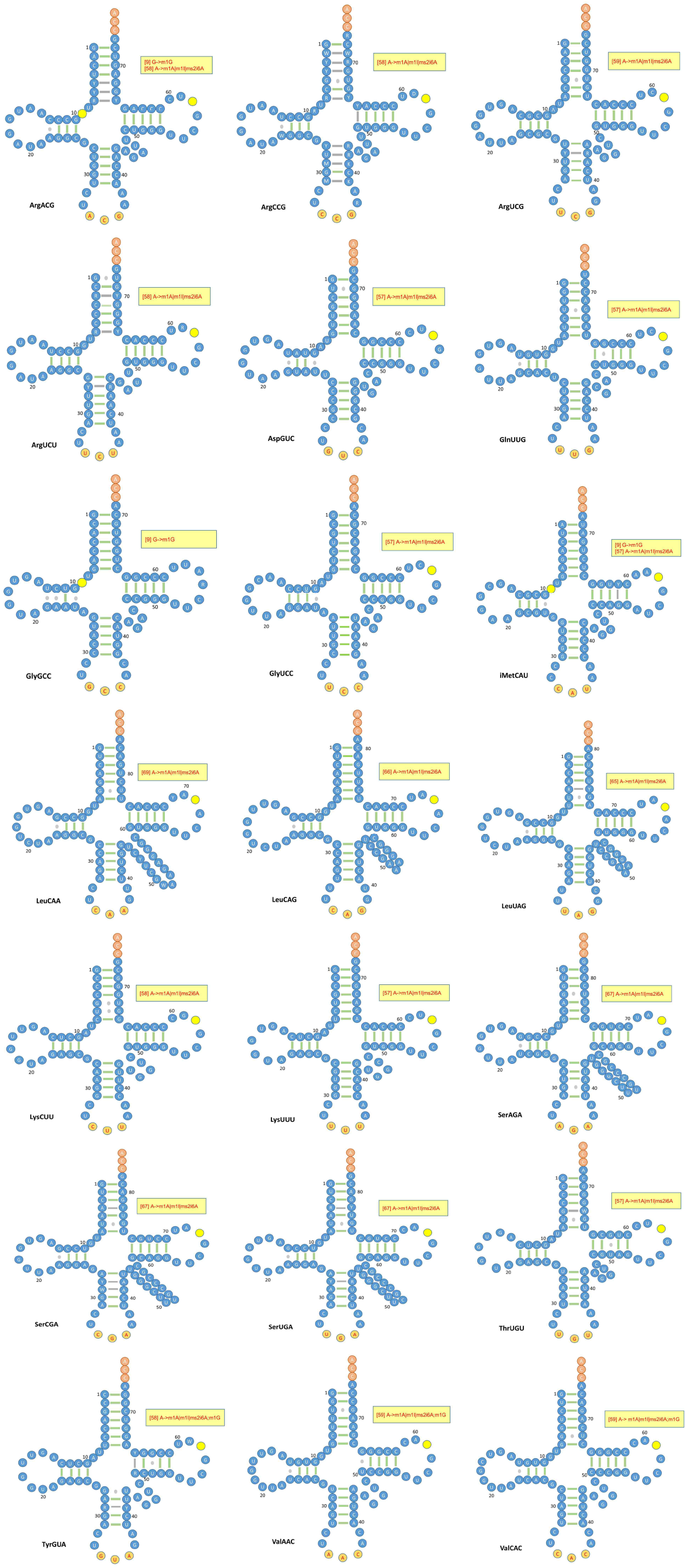
tRNA isoacceptor secondary structure models showing common nucleoside modifications in all six plants. Circles (in yellow) depict the position of modification residue on the tncRNA generated from the respective tRNA.

### 2.6. tncRNAs are conserved amongst plants and can play a paramount role as riboregulators

In our total analysis, we observed various tncRNAs were identical in all six plants. We found a total of 252 tRF-5, 351 tRF-3, 46 5’tRH, and 12 3’tRH common sequences. The sequence conservation provides a clue to the conservation in the regulatory function of these tncRNAs. tncRNAs can act as post-transcriptional regulators by complementary binding to their messenger RNA (mRNA) targets leading to mRNA degradation. Thus, for gaining some functional insights, some of these conserved sequences were utilized for target prediction and pathway enrichment analysis. The detailed list of conserved tRF-5s and tRF-3s along with their respective target genes have been provided in Supplementary sheet 2.1 and 2.2 respectively. We observed that several target genes were associated according to the molecular function, biological process, and cellular components for tRF-5 and tRF-3. The tRF-5 target genes were related to 9 distinct molecular functions, 25 types of biological processes, and 6 cellular components (Fig. 6A). The tRF-3 target genes were associated with 13 molecular functional categories, 26 biological activities, and 7 cellular components (Fig. 6B). The target genes for the tRF-5 series were involved in various molecular functions such as transferase, catalytic, kinase, endopeptidase, actin-dependent ATPase, and microfilament motor activities. Genes were enriched in various metabolic processes, transport, localization, and developmental processes. The target genes for the tRF-3 series were associated with numerous activities like catalytic, transferase, ATPase, nucleoside-triphosphatase, RNA helicase, etc. The list of all molecular functions, biological processes, and cellular components along with their Gene Ontology (GO) term ID, Padj values, and −log_10_(Padj) for predicted tRF-5 and tRF-3 targets is shown in Fig. 6C and Fig. 6D respectively. A detailed list for tRF-5 and tRF-3 GO has been provided in Supplementary sheets 2.3 and 2.4 respectively. For a more organized visualization of our enrichment results, we clustered our enriched genes into enrichment maps. Clusters of similar pathways representing major biological processes were generated. The tRF-5 target gene clusters were enriched in growth and development, transport and localization, and cellular metabolic processes were observed (Fig. 6E). tRF-3 target genes formed two distinct clusters related to developmental and cellular metabolic processes (Fig. 6F). It can be assumed that many tRF-5 and tRF-3 may act as potent regulators of the genes involved in plant development and cellular metabolic machinery in a tissue/condition-specific manner. Although many studies have shown that tRFs modulate gene expression [43], [66], [67] i.e. by binding to mRNA (UTRs or exon regions) in a sequence complementary manner and direct transcript cleavage, we have seen that a single tRF may target multiple transcripts. Although how a single tRF can regulate multiple target transcripts is still unclear to the researchers.

**Figure 6:**
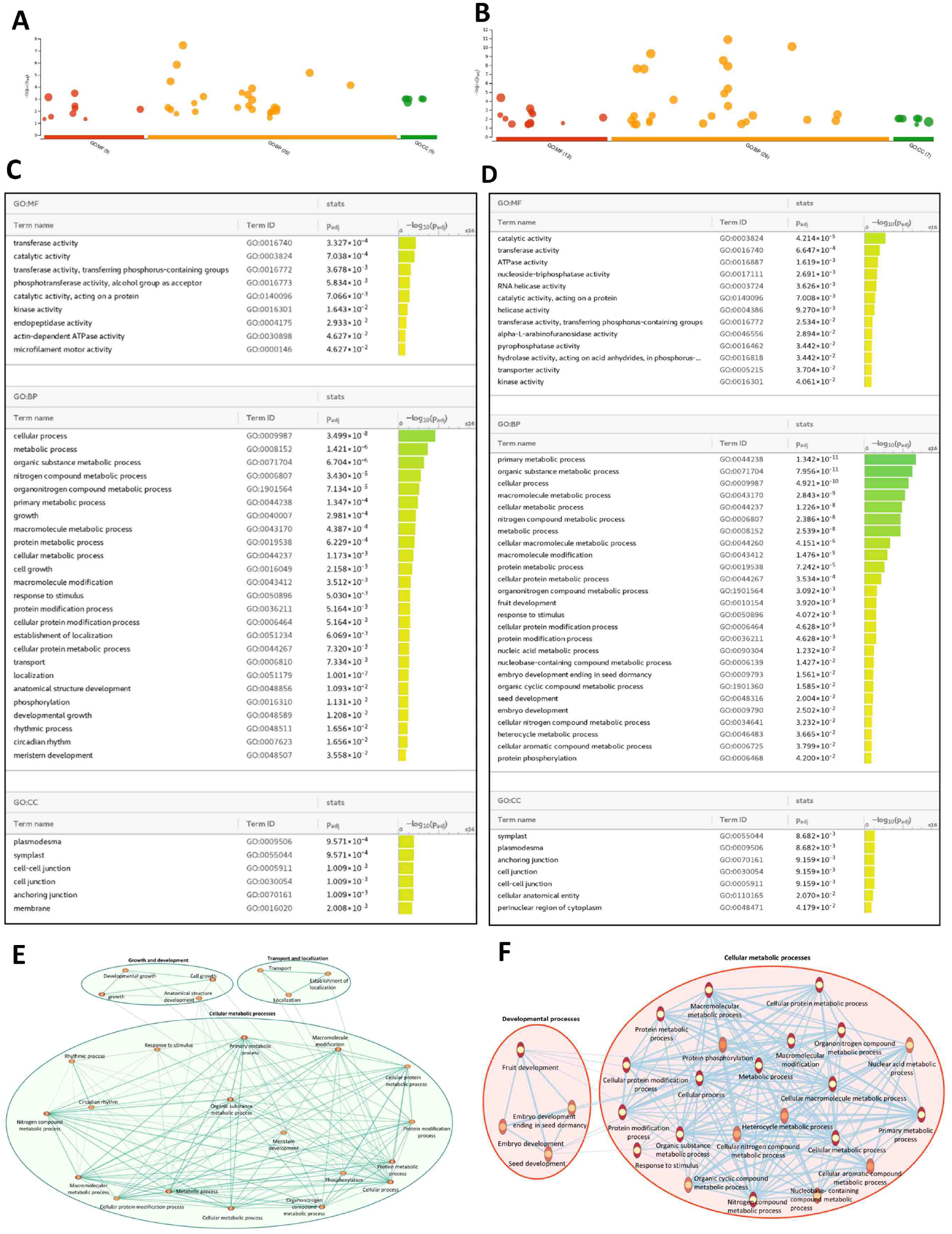
Pathway Enrichment analysis for conserved tRF-5 and tRF-3 target genes. **A,B** Manhattan plot illustrating the enriched terms across the analyzed GO term categories for tRF-5 and tRF-3 targets respectively. The x-axis shows the functional terms and the corresponding enrichment P-values in the negative log10 scale are represented on the y-axis. Each color-coded circle on the plot represents a single functional term. **C,D** Detailed results in tabular output illustrating GO term name, term ID, and P-values (in order of the most to least significant) for tRF-5 and tRF-3 targets respectively. **E,F** Enrichment map visualization of enriched pathways generated using. **G:**Profiler analysis for tRF-5 and tRF-3 target genes respectively. Pathways are shown as circles (nodes) that are connected with lines (edges) if the pathways share multiple genes. Nodes and edges were manually formatted to present a neat picture.

We also examined conserved tRF-5 and tRF-3 against full transposons elements (TEs) from *Arabidopsis*. We observed that many tRF-5 and tRF-3 sequences were associated with several TE transcripts (Supplementary sheets 2.5 and 2.6). A large number of tRF-5 showed association with various transposon families (Fig. 7). As compared to tRF-5s, tRF-3 showed a weaker association with TEs. TE belonging to the LTR/Gypsy superfamily were the most frequent targets of the tRF-5 series. With this superfamily, TE from *ATHILA2, ATHILA6A, ATHILA4C, ATHILA4A, ATLANTYS1, ATHILA6B*, and *ATLANTYS2* families were found to be associated with a large number of tRFs. Apart from them, TE superfamilies like RC/Helitron (*ATREP3* and *ATREP15*) and DNA/Mudr (*VANDAL2* and *VANDAL3*) also show considerable association with tRFs. Smaller length tRFs (mostly 19-22 nt) originating from AspGTC/ATC, GlnTTG, ProAGG/CGG/TGG, HisGTG, LeuCAA/TAG, and SerGCT tRNAs were associated with TE elements in high numbers. Binding to TE by tncRNAs can bestow genome stability to plants. Interestingly, two recent reports have shown that tRFs target transposable elements (retrotransposons) in both plants and mammals, indicating their potential as epigenetic regulators [68]. E.g., in *Arabidopsis*, a 19 nt tRF-5 from tRNA-MetCAT cleaves its target LTR retrotransposon transcript, *Athila6A* [69]. Deciphering the synchronization between the tncRNA generation, TE expression, and their implications in plants under normal and stress conditions can unveil additional regulatory functions of this class of ncRNAs.

**Figure 7:**
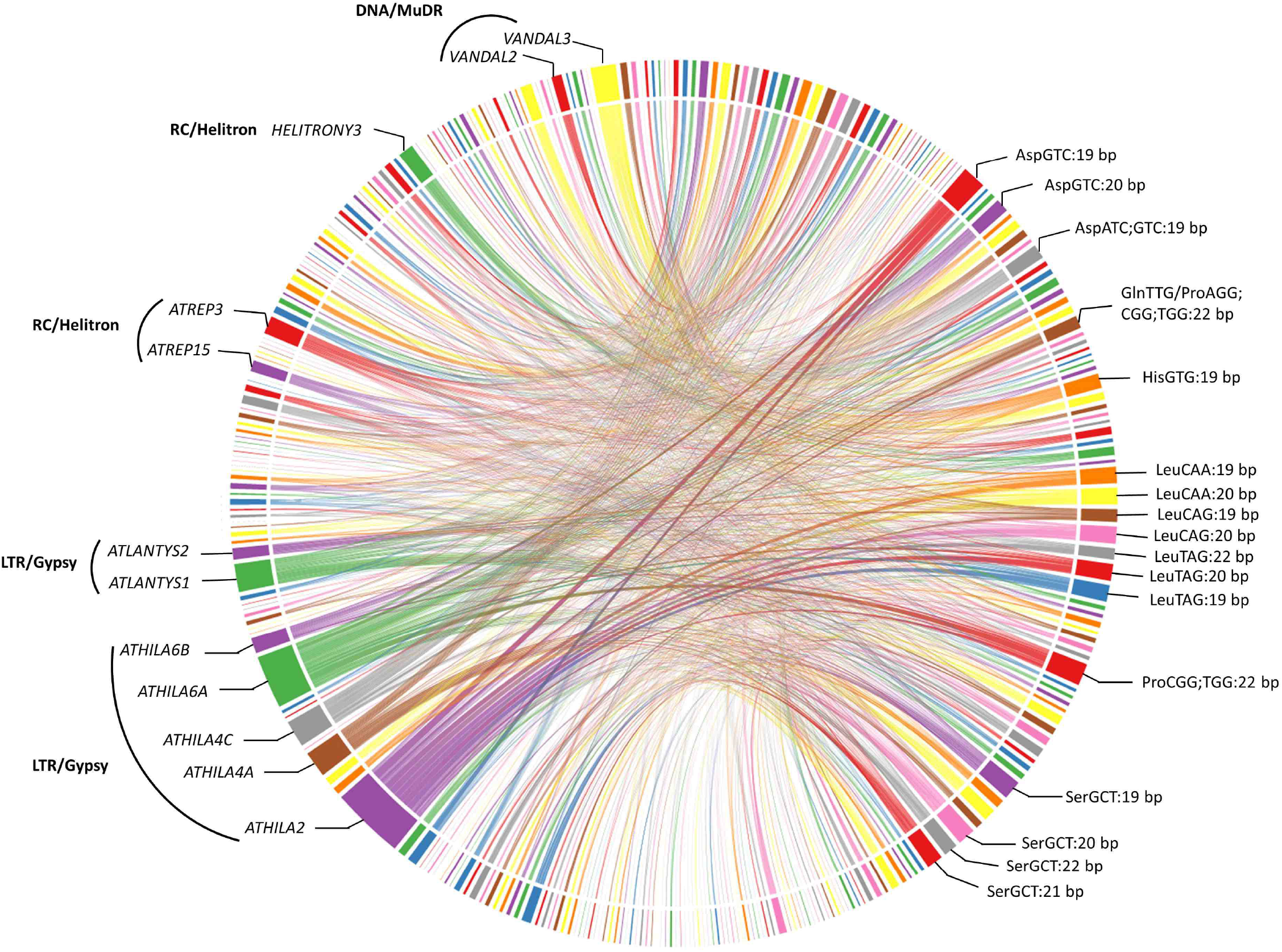
A chord diagram representing the connections between conserved tRF-5s and their corresponding target transposable elements (TE) family and their superfamily. Most recurring TE family and associated tRF-5 have been labelled. Broader fragment reflects the greater number of associations.

### 2.7. Identification of tissue-specific tncRNAs from *Arabidopsis*

A total of 525 small RNA-seq samples (Supplementary sheet 3.1) of *Arabidopsis* were analyzed for the identification of tissue-specific tncRNAs. To determine whether tncRNAs show specificity towards distinct tissues in plants, we used t-Distributed Stochastic Neighbour Embedding (t-SNE), a dimensionality reduction technique to graphically simplify and explore very large datasets. We utilized this technique to explore tncRNA heterogeneity in 525 SRA samples covering 13 major tissues from the *Arabidopsis* plant. t-SNE plots revealed numerous tissue-specific clusters for individual tncRNA classes (Fig. 8). Seedlings and seed clusters were frequently seen in tncRNAs belonging to tRF-5, tRF-3, tRF-1, and 5’tRH. As compared to the tRNA halves, smaller fragments particularly belonging to tRF-5, and tRF-3 series showed enriched clustering for more than ten different tissues. Very few clusters were seen for 5’tRHs originating from seedlings, seed, inflorescence, silique, flower, etc. It was interesting to observe that vasculature and epidermal tissue clusters were found to be in close proximity in tRF-5, tRF-3, and 5’tRHs. Surprisingly, seedling, seed, shoot, rosette leaf, inflorescence, silique, and flower clusters were also seen in the tRF-1 series. The pattern of distribution of tncRNAs belonging to various tissues in *Arabidopsis* indicates that spatiotemporal expression of tncRNAs occurs in a tissue-specific manner at normal physiological conditions. Tissue-specific tncRNA signatures might regulate tissue differentiation, organ development, and regeneration in plants.

**Figure 8:**
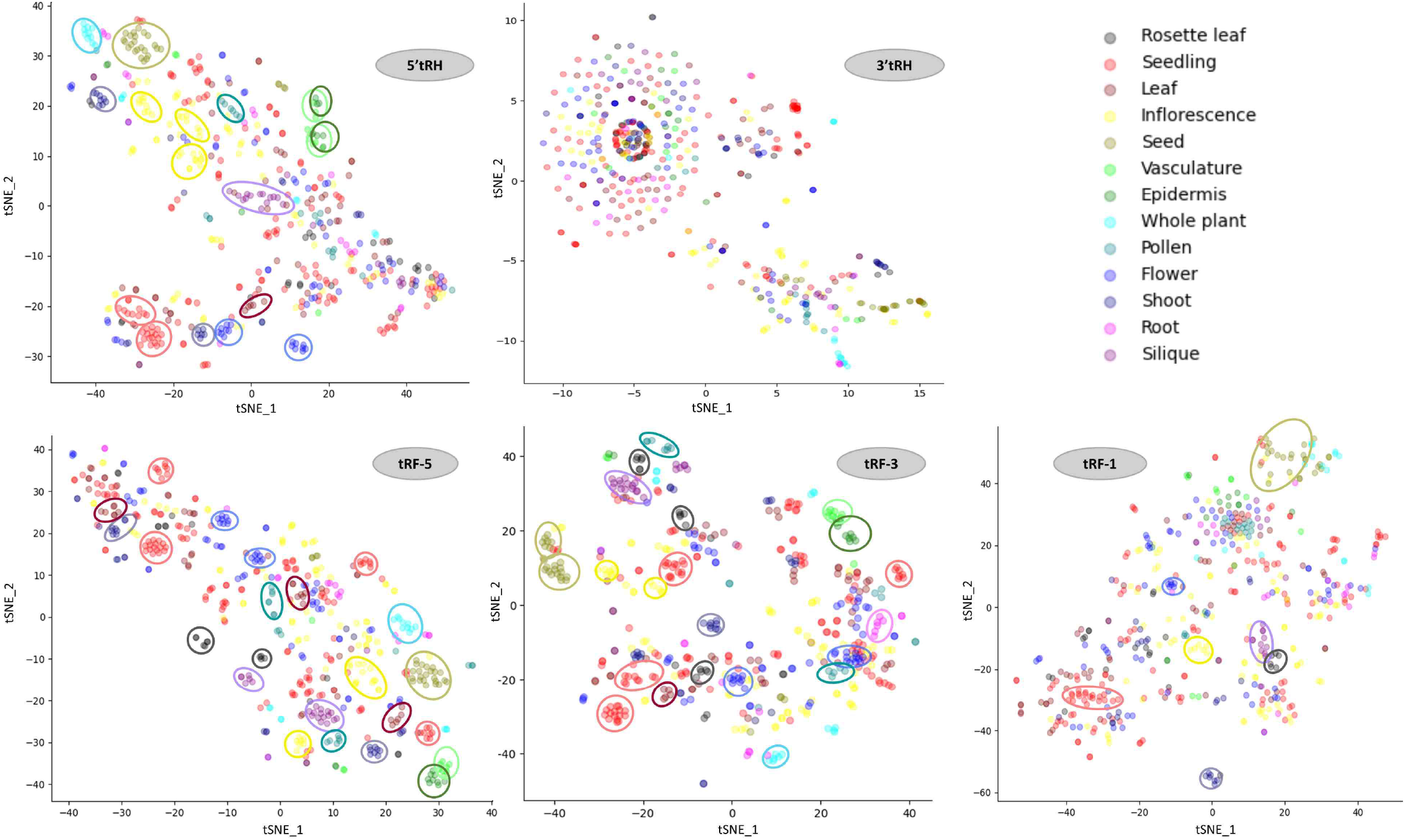
t-Distributed Stochastic Neighbor Embedding (t-SNE) for the visualization of the tissue-specific tncRNA expression in *A. thaliana* (wild type under normal conditions) datasets. Individual t-SNE plot was made for tRF-5, tRF-3, tRF-1, 5’tRH, and 3’tRH classes. The tncRNAs from a total of 525 samples from 13 different tissues were utilized for this analysis. Tissue clusters observed have been shown by similar color-coded dashed circles.

### 2.8. Different classes of tncRNAs are differentially expressed (DE) under various conditions

To calculate the expression of tncRNAs under various abiotic and biotic stresses, a total of 168 small RNA-seq samples viz. *Arabidopsis* (57), rice (36), tomato (6), and maize (69) were analyzed (Supplementary sheet 3.2) We checked the expression of tncRNAs under various abiotic and biotic stresses in *Arabidopsis*, tomato, rice, and maize (Supplementary sheet 3.3-3.6). Significant differences in tncRNA expression were observed in different plants under different conditions. Besides up-regulated fragments, a significant number of tncRNA were down-regulated in various samples. The DE tncRNAs expressed in more than one sample under different conditions in *Arabidopsis*, and in at least five samples in maize have been shown in Fig. 9A and 9B respectively.

**Figure 9:**
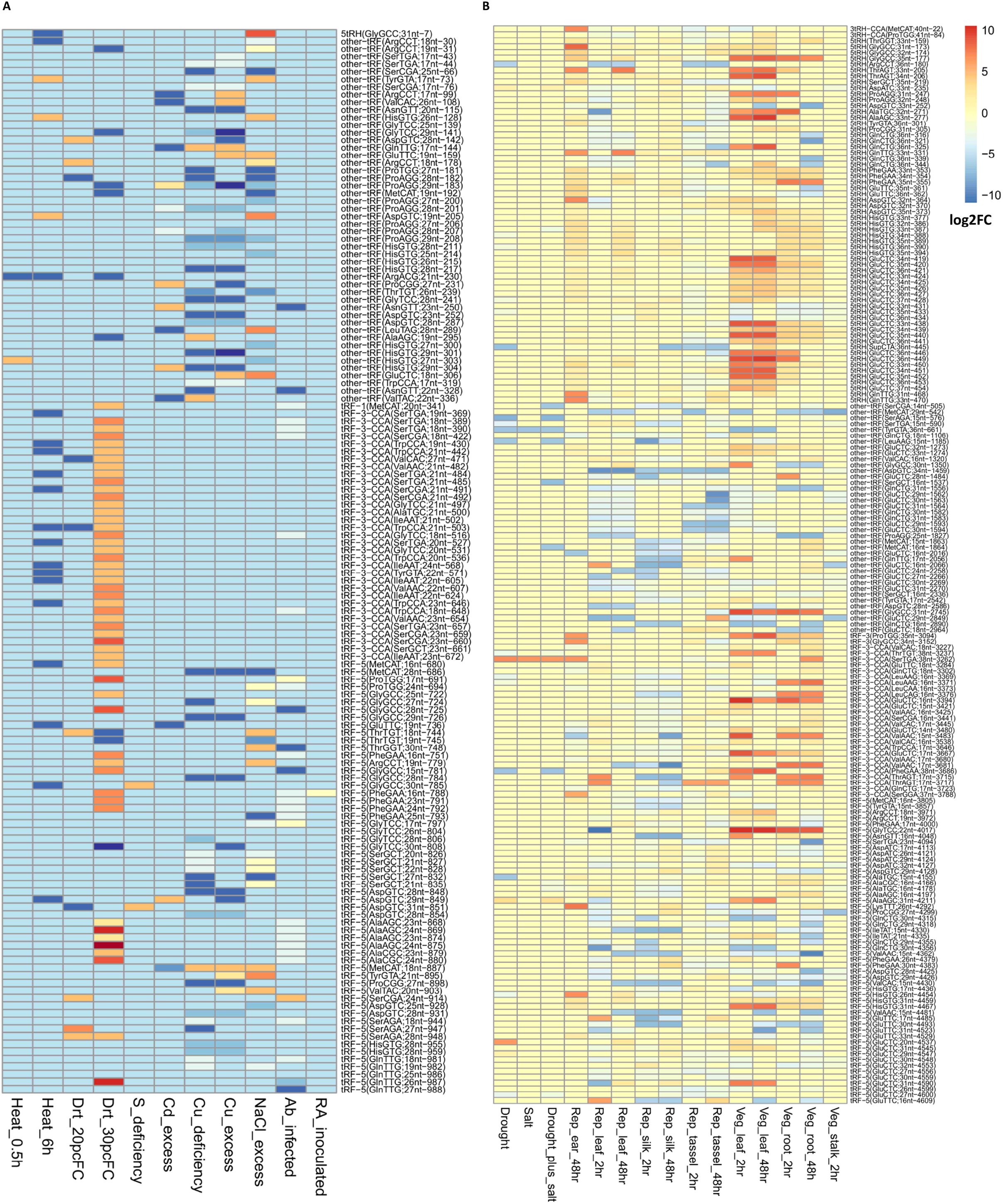
Visualization of differentially expressed (DE) tncRNAs in *A. thaliana* and *Z. mays* (maize). Heatmap was built using the log2FC values, in the color scale, red represents positive log_2_ fold-change (log_2_FC) and blue represents negative log_2_FC. **A** DE expressed tncRNAs in *A. thaliana* in eleven different types of stress from left to right viz. heat stress (0.5hr), heat stress (6hr), drought stress (at 20% field capacity), drought stress (at 30% field capacity), sulfur deficiency, cadmium excess, copper deficiency, copper excess, NaCl excess, *Alternaria brassicicola* infection, and Rhizobacteria inoculated samples. **B** DE expressed tncRNAs in Z. mays in fifteen different conditions with respect to stress condition and tissue origin viz. drought, salt, drought plus salt, heat stress (2hr, 48hr) in various reproductive and vegetative tissues. To reduce the large number of DE tncRNAs detected in various samples, we have represented only those tncRNAs in the heatmap occurring in at least 2 samples in Arabidopsis and five or more samples in maize. The full list DE tncRNAs for *Arabidopsis* and maize has been provided in Supplementary sheet 3.3 and 3.6 respectively.

We observed stress, time, or tissue-dependent tncRNA expression in different plants. In *Arabidopsis*, only 3 different tncRNAs showed dysregulation after 0.5h of heat stress but their number rose to 42 after a time duration of 6 h. Likewise only 25 tncRNA sequences were expressed drought stress at 20% field capacity (FC). However, 182 DE tncRNAs were detected when FC was 30% FC drought stress. Less than 10 DE tncRNAs were detected in *M. oryzae* infected root tissues (9 and 7 DE tncRNAs after 30 and 120 min respectively) (Fig. 9A). However, the numbers were highly elevated in infected leaves as compared to root tissues (87 and 62 tncRNAs after 30 and 120 min respectively). The time-dependent difference in expression was also observed in few tissues in maize under heat stress (2h vs 48h). In RA-inoculated *Arabidopsis* root tissues, 212 unique tncRNA sequences were differentially expressed.

Among all the conditions studied, we found that heat exposure leads to the maximum number of differentially expressed tncRNAs in different tissues of maize, both in vegetative and reproductive stages when compared with control. The highest among all with a total of 1748 tncRNAs were differentially expressed in tassel in reproductive stage upon exposure to 48 hours of heat stress, among which 35 nt 5’tRH derived from tRNA-GlyTCC, was significantly downregulated with a log_2_FC of −30. Of the 469 DE tncRNAs in *Arabidopsis* upon *Alternaria brassicicola* infection, 266 were upregulated and 203 were downregulated. Besides, Rhizobia inoculated sample, drought (30%FC) and excess NaCl conditions in *Arabidopsis* revealed a considerable number of DE tncRNAs. Of the 989 DE tncRNAs that were expressed in at least one stress condition in *Arabidopsis*, mostly ranged from 17-27nt, with an abundance of 18 nt (125) followed by 17 nt (86), and 19 nt (79) long tncRNAs. The tomato plant when subjected to Tomato Mosaic Virus (TMV) infection showed 757 DE tncRNAs, of which the majority of the fraction were contributed by 15 nt tRFs. While the total abundance of up- and down-regulated tRFs were majorly contributed by other tRFs, tRF-5 and tRF-3(CCA) were the conventionally known major tncRNAs that played an important role in different stress responses. Further, certain tRFs were found to be overexpressed or significantly downregulated and thus followed an unusual trend unlike the tRFs from the same population; for instance, two tRF-5s (GlyTCC;27 nt and AlaAGC;24 nt significantly upregulated at 30%FC drought stress in *Arabidopsis* with log_2_FC of 27.71 and 27.62 respectively, whereas a 17 nt tRF-5 derived from ValCAC was highly upregulated upon inoculation of *Alternaria brassicicola* with a log_2_FC of 20.88. Also, it was noticed that DE tncRNAs in *Arabidopsis* were generated from HisGTG at a comparatively elevated level than others, GlnCTG in maize, GluTTC in rice, and tomato.

We also observed DE tncRNAs belonging to the same class generating from the same tRNA. As an example, two 19 mer tRF-5s from tRNA-LeuCAA (GTCAGGATGGCCGAGTGGT and GTCAGGTTGGCCGAGTGGT) differing only at one nucleotide at the 7th position was found to be up-regulated under NaCl stress. Identical tRF-5s derived from tRNA-AspGTC of 14, 16, 17, and 19 nt were all equally upregulated in RA-inoculated *Arabidopsis* samples. This indicates that similar tncRNAs of varying length may evoke a similar response and function complementarily under particular conditions. Surprisingly, some DE tRF-1 fragments were also detected in *Arabidopsis*, tomato, rice, and maize (Supplementary sheet 3.3-3.6). A 20 nt tRF-1 derived from tRNA-MetCAT was accumulated under drought as well as *A. brassicicola* infected *Arabidopsis* plants. In our results, we also observed the upregulation of 19 nt tRF-5 from tRNA-ArgCCT under drought (30%FC) and NaCl stress in *Arabidopsis*. The same tRF also showed elevated expression in maize leaf subjected to heat stress. In earlier studies, it was reported that the same tRF was expressed at high levels in *Arabidopsis* seedlings under drought conditions as well as in cold-treated rice inflorescences [34]. The findings suggest that the accumulation of evolutionarily conserved plant tncRNAs is regulated by biotic and environmental stresses. Further, validation of DE tncRNAs and their putative targets will pave a way for a better understanding of the underlying mechanism in the regulation of stress responses.

## 3. Discussion

To the best of our efforts, we have designed a workflow to accurately identify genuine tncRNAs from small RNA-seq datasets. We have developed our workflow for convenient tncRNA mining and can be tailored as per the requirement. Considering various computational limitations in mapping tRNA fragments, we have tried to eliminate false positive and ambiguous fragments from our analysis using the computational pipeline. Nevertheless, RNAs of such a small size range make it difficult to distinguish them from randomly cleaved degraded products. Genome-scale study of tncRNA expression in angiosperms revealed that there are various classes of tncRNAs generated from distinct portions of genomic tRNAs (Fig. 10). tRNA gene clusters are localized at variable intervals over the genome and they may also affect tncRNA generation. For example, tRNA-dense regions can be seen particularly on *Arabidopsis* (chromosome 1), tomato (chromosomes 1 and 6), chickpea (chromosomes 1 and 5), *Medicago* (chromosomes 1 and 4), rice (chromosomes 4 and 10), and maize (chromosomes 4 and 5). However, it can be observed that the tncRNAs are not necessarily generated from tRNA-rich regions of the genome. Also, all classes of tncRNAs are generated from distinct genomic regions in different plants. Interestingly some regions also produce most classes of tncRNAs, like on chromosome 2 (*Arabidopsis*), chromosome 1 (tomato), chromosome 4 (*Medicago* and rice), and chromosome 5 (maize). Some regions were producing more 5’ containing tncRNAs i.e. tRF-5 and 5’tRHs, on chromosome 3 in *Arabidopsis*, chromosome 6 in tomato, chromosome 4 (*Medicago*), chromosome 11 (rice), and chromosome 5 (maize). It is quite conceivable that some of these regions may govern rapid turn-over of tRNAs into tncRNAs. All of these findings giving clues about the additional insights into the genomic organization of tRNA into clusters and their implications on tRNA functions. The prevalence of tRNA gene clusters in all kingdoms of life reveals various insights into evolutionary history and tRNA functions. Whether tRNA gene clusters impact the generation of tncRNAs in the living system including plants is worth intriguing.

**Figure 10:**
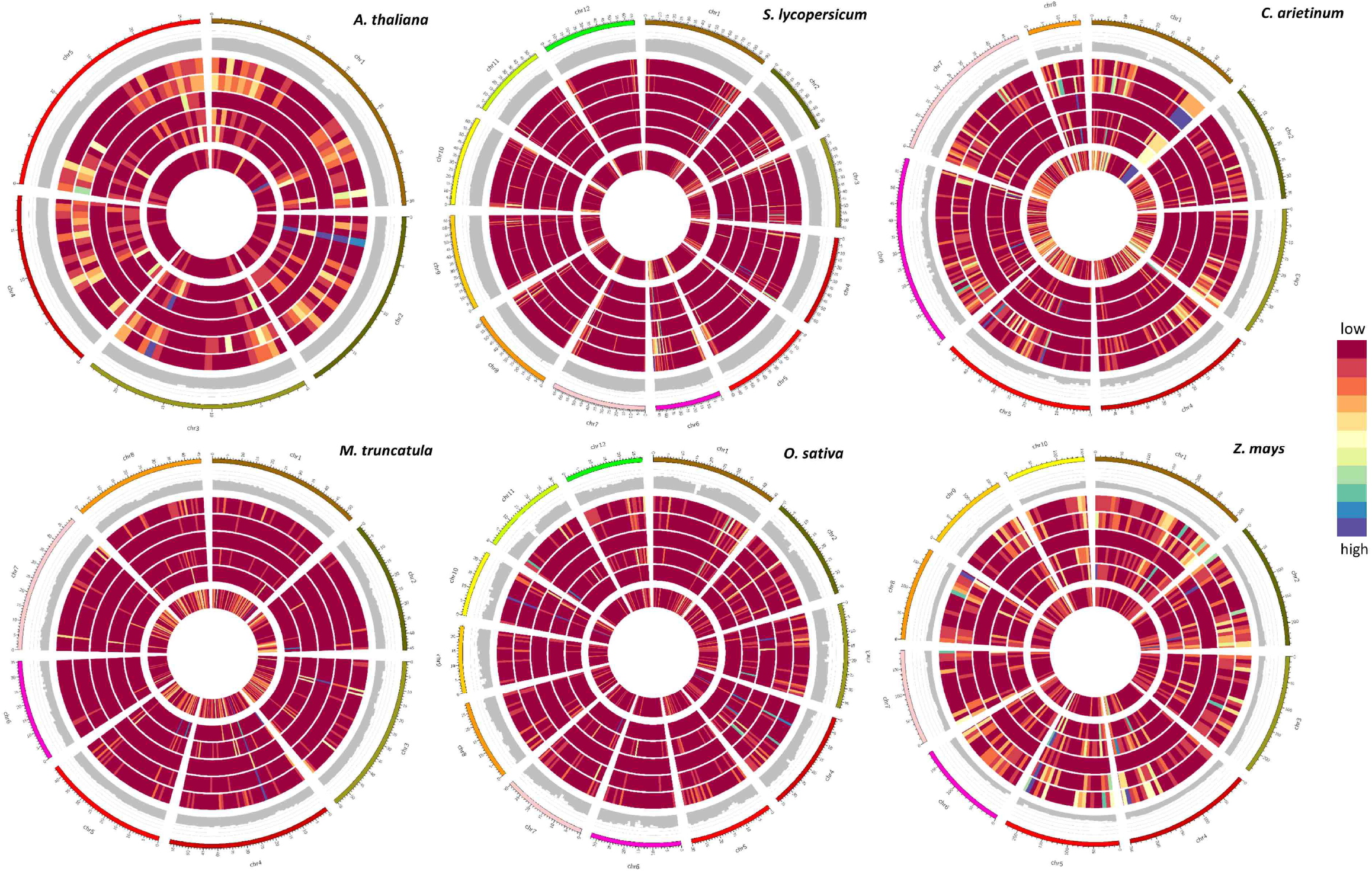
Circos plots for interactive visualization of the genomic origin of various tncRNAs subclasses generated from parental tRNAs and the relative position of tRNA genes on the genome for individual plants. The relative abundance of various tncRNA subtypes and tRNA gene density with respect to the genomic location is shown by the heatmap. From the outer to inner circle, the first ring represents the chromosomes, the next ring (second) shows the GC content of the chromosomes. The third, fourth, fifth, sixth, and seventh ring shows the genomic tRNA region producing tRF-5, tRF-3(CCA), tRF-1, 5’ tRH, and 3’tRH respectively. The innermost ring shows the tRNA gene distribution across the genome. tncRNA/tRNA-rich regions shown in purple (highest density) and tncRNA/tRNA-poor regions shown in the red (minimum density) spectrum.

In plants, tncRNAs can be also generated from two more organelles besides the nucleus. In addition to the nucleus, the chloroplast and mitochondria also generate tncRNAs. Excluding those tncRNAs that can be mapped on multiple locations, a considerable number of tncRNAs originate exclusively from either nucleus, chloroplast, or mitochondria. For example, in *Arabidopsis*, 24% tRF-5s and 33% tRF-3s originated from either mitochondria or chloroplast. Among tRHs, 29% 5’tRHs and 25% 3’tRHs were exclusively derived from organellar tRNAs. Overall among different plants, tncRNAs generated from plastid are more in number as compared to mitochondria. However, in maize datasets, most of the organellar-derived tncRNAs were mitochondrial in origin. The counts of exclusive unique tRF-5, tRF-3, 5’tRH, and 3’tRH separately for all plants have been provided in Supplementary sheet 1.7. The complex pool containing mitochondrial, and plastidial encoded tncRNAs detected in our datasets indicate the hidden machinery driving their biogenesis, transport in the cytosol, and their functions in the cellular milieu. Although some organellar-derived tncRNAs have been shown to accumulate in the cytoplasm [70], how they are generated, transported outside the organelle and their functions remain to be elucidated.

tncRNAs particularly in the length range of 19-25 nt have been studied for the AGO interaction and AGO-IP libraries have been observed to be rich in tRNA fragments [33], [34]. For instance, in our tissue as well as stress-related studies we have seen the abundance of tncRNAs in non-AGO IP libraries as well. This also indicates that AGOs may or may not associate with all tncRNAs. Apart from conventional AGO binding mediated RNA interference, many tncRNAs may be functionally independent of AGO, unlike miRNAs. The expression of tncRNAs in non-AGO IPs indicates some other mechanisms might also in fine-tuning gene expression. The production of tncRNAs from precursor tRNAs has not been well studied in plants. Till now in plants, tRF-1 fragments have been rarely identified and characterized. In humans, tRF-1/pre-tRF-3U usually end in a short stretch of T residues due to the release of polymerase III[6], [71]. However, in plants, most but not all tRF-1 sequences have a stretch of poly(T) residues in the terminal end (Supplementary Figure 10). The majority of tRF-1 sequences identified in rice, maize, and tomato samples had terminal end rich in T residues. Interestingly, all four DE tRF-1 in tomato infected with tomato virus and short DE tRF-1 (18-28 nt) in maize ended in poly(T) tail (Supplementary sheet 3.4 and 3.6). Although some general mechanisms appear to be universal and conserved across kingdoms, there exist additional determinants that specifically drive the biogenesis of different tRNA-derived fragments in plants. Also, the DE tRF-1 reported in plants for the first time in our study encourages us to commence the exploration of these fragments *in planta*.

The identification of genuine tncRNAs is just a head start to explore innumerable pathways that are involved in the cellular machinery and to investigate their putative targets. Although the *in-silico* approach will help in identifying these novel molecules across a wide range of organisms, experimental validation, and characterization are vital for distinguishing more and more *bona fide* tncRNAs from randomly degraded tRNA fragments. tncRNAs play biological roles through a variety of mechanisms by interacting with proteins or mRNA, inhibiting translation, and regulating gene expression, the cell cycle, and chromatin and epigenetic modifications. Switching from conventional RNA-seq to tRNA-seq to overcome modification biases will be advantageous for the identification of tncRNA. To study the molecular mechanisms of tncRNAs, apart from classical molecular biology methods, such as northern blotting, microarray, real-time quantitative and reverse transcription-polymerase chain reaction (qRT-PCR); advanced ribonomics [72] like cross-linking, ligation, and sequencing of hybrids (CLASH); photoactivatable-ribonucleoside-enhanced cross-linking and immunoprecipitation (PAR-CLIP) may lay the foundation for studying tncRNA-target hybrid in plants. As tncRNAs are ubiquitous and conserved across different domains of life, our comprehensive study is believed to substantiate research of these novel molecules. This will aid in the investigation of non-canonical functions of tRNA, the critical role of tRNA modifications in tncRNA genesis, and implications of tncRNAs in biological pathways governing cellular physiology under different stress conditions, tissues, and developmental stages. Their global identification will facilitate deciphering common conserved pathways in eukaryotes and their mechanism of unexplored regulatory action. Leveraging computational power together with molecular biology techniques will augment the current understanding of the vast tncRNAomic landscape in the living system.

## 4. Methods

### 4.1. Data retrieval

Genome assembly for *A. thaliana* (GCF_000001735.4), *S. lycopersicum* (nuclear: solgenomics build_4.00; organellar: GCF_000188115.4), *C. arietinum* (GCA_000331145.1), *M. truncatula* (nuclear: GCF_000219495.3; organellar: NC_029641.1), *O. sativa* (GCF_000005425.2), and *Z. mays* (GCF_902167145.1) were downloaded, and IDs were transformed for starting with “chr”, to make them convenient for secondary analysis.

Single-end small RNA-Seq datasets (Illumina) available for the abovementioned plants were downloaded from NCBI SRA (https://www.ncbi.nlm.nih.gov/sra) using SRAToolkit (v2.10).

### 4.2. tRNA annotation and genome pre-processing

Nuclear tRNAs were predicted by tRNAscan-SE [60] (v2.0.6) using the eukaryotes model, while organellar tRNAs were detected by using the ‘-O’ option in this tool. The tRNA pseudogenes and tRNAs with a score less than 50 were eliminated, and filtered tRNA regions with 50 base pair (bp) flank at 3’ and 5’ were masked in the reference genome by ‘maskfasta’ script of bedtools [73] (v2.29). Further, three FASTA files were prepared, 1) mature tRNA, built from filtered tRNAs after intron removal, and addition of CCA at 3’ terminus, 2) leader tRNA, 50 bp 5’ tRNA genomic region flank, and 3) trailer tRNA, 50 bp 3’ tRNA genomic region flank. These FASTA files were added to the masked genome as ‘additional chromosomes’ to create an artificial genome. The Bowtie index for the artificial genome was built by bowtie-build (bowtie v1.3).

### 4.3. Identification of modification site

The single-end small RNA reads were processed by TrimGalore (v0.6.6) to remove the low-quality bases, and to trim the adapter sequences. To permit error-tolerant, and uniquely mapping reads alignments, processed reads were aligned to the artificial genome index using bowtie[74] with ‘-v2 --best’ options. The SAM output was sorted and converted to BAM by samtools [75] (v1.10). The modification sites were predicted by HAMR [64]. Further, modification sites only on mature tRNA and leader/trailer regions were only counted.

### 4.4. tncRNAs identification and classification

The mapped reads from previously generated SAM files were fetched, to create a FASTA file of unique reads with the count, using SAM flag 0 & 16 for single-end reads. Mapped unique reads were aligned to +ve strand only with no mismatch, and multi-mapped reads were discarded if alignment occurred >50 times, using bowtie1 with “--norc -v 0 -m 50” arguments. The bowtie combinatorial arguments “--best” and “--strata” were used which guarantees that reported singleton alignments are best in terms of the stratum. Then, the output was used to identify locus, location, length of read mapped to mature tRNA, and their 50 bp upstream and downstream flank; based on that, reads were classified into different classes of tncRNAs. The tRNA halves were classified by cleavage at the anticodon loop (2 nt + 3 nt of anticodon + 2 nt = 7 nt), and information for modification site was added for tncRNAs. Reads per million (RPM) was calculated for each tncRNA by using the formula:

Per Million Factor (PMF) = Total mapped reads /10^6^

Reads per million (RPM) = Number of reads mapped to progenitor tRNA / PMF

A local alignment file of tncRNA to the sequence of origin was also created for visualization by using the pairwise2 biopython module. The alignment score is given as per

*identical=1, non-Identical= −1, gap-open= −1, gap-extend= −0.5*

### 4.5. tRNA model for consensus sequence and structure

The FASTA files were generated for specific isoacceptor tRNAs and provided as an input to the LocARNA [76] (v1.9.2) software package with “--stockholm” option for consensus sequence study. These ‘stockholm’ format files were used to create postscript files by RNAalifold [77] (v2.4.11) for the consensus tRNA sequence. The modification site(s) for each tRNA isoacceptor has been shown on the consensus tRNA structures.

### 4.6. Target prediction, GO, and pathway enrichment

The psRNATarget [78] software was utilized for tncRNA target prediction (2017 release; default parameters). Common tncRNAs (tRF-5 and tRF-3 series; 18-25 nt long) were used as a query while *Arabidopsis* cDNA and transposons sequences were used as probable targets. The protein targets were analyzed for pathway enrichment analysis using g:Profiler[79] and visualized in Cytoscape [80] (EnrichmentMap and AutoAnnotate).

### 4.7. t-Distributed Stochastic Neighbor Embedding (t-SNE) plot

t-SNE is a non-linear technique for dimensionality reduction that is particularly well suited for the visualization of high-dimensional datasets. The samples sourcing from wild-type *Arabidopsis* (Col-0) grown under normal conditions belonging to different tissues were selected for this study. A total of 525 samples were utilized belonging to 13 unique tissues viz. epidermis, flower, inflorescence, leaf, pollen, root, rosette leaf, seed, seedling, shoot, silique, vasculature, and whole plant. A binary matrix, for presence (1) and absence (0), was created for unique tncRNA sequences from each tissue sample. From this, a t-Distributed Stochastic Neighbor Embedding (t-SNE) plot was generated for each aforementioned class of tncRNAs. Sample and tissue information for this plot is provided in Supplementary sheet 3.1.

### 4.8. Differential expression study

DESeq2 was utilized for the identification of differentially expressed tncRNAs under stress conditions. Only those samples were selected in which at least three biological replicates for each control and treatment were present (as provided in Supplementary sheet 3.2). In DESeq2, the *p*-values attained by the Wald test are corrected for multiple testing using the Benjamini and Hochberg method by default [81]. For our analysis, only those tncRNAs were considered to be significant with *P* less than 0.05. Only significant tncRNAs were utilized for expression analysis. tncRNAs with log_2_FC value greater than 1 (>1), and lesser than −1 (<-1) were considered up- and down-regulated respectively. The ‘pheatmap’ package of R was used to draw the heatmaps of significant differentially expressed tncRNAs.

## Supporting information

Supplementary sheet 1

Supplementary sheet 2

Supplementary sheet 3

## Code and Data availability

All pipeline scripts, codes, data generated, and analyzed for each of the species are available at the tncRNA website (URL: http://nipgr.ac.in/tncRNA).

## Author contributions

S.Z. and A.S. designed the pipeline, wrote the codes, and performed all the analysis. N.P. contributed to the data analysis. S.Z. and SK wrote the manuscript. SK conceived the study and coordinated the project.

## Acknowledgments

S.Z. and A.S. acknowledge the Council of Scientific and Industrial Research (CSIR), India, for Senior Research Fellowship. This study was supported by Grant SRG/2019/000097 from Science and Engineering Research Board (SERB), Department of Science and Technology, Government of India. The authors are thankful to the Department of Biotechnology (DBT)-eLibrary Consortium, India, for providing access to e-resources.

