## Supplementary data for "Transfer RNA-derived non-coding RNAs (tncRNAs): Uncovering hidden regulators of transcriptional regulatory circuits in plants"

**Supplementary figures**

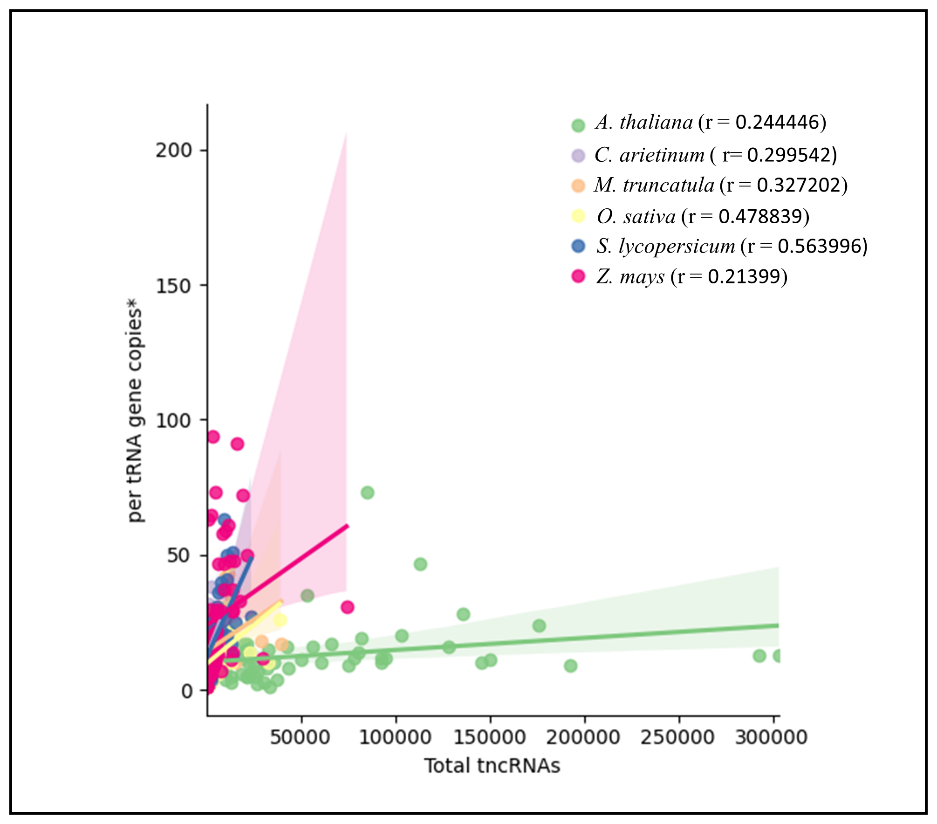

Supplementary Figure 1: Correlation graph showing the total tncRNAs identified vs per tRNA gene copies in individual plants.

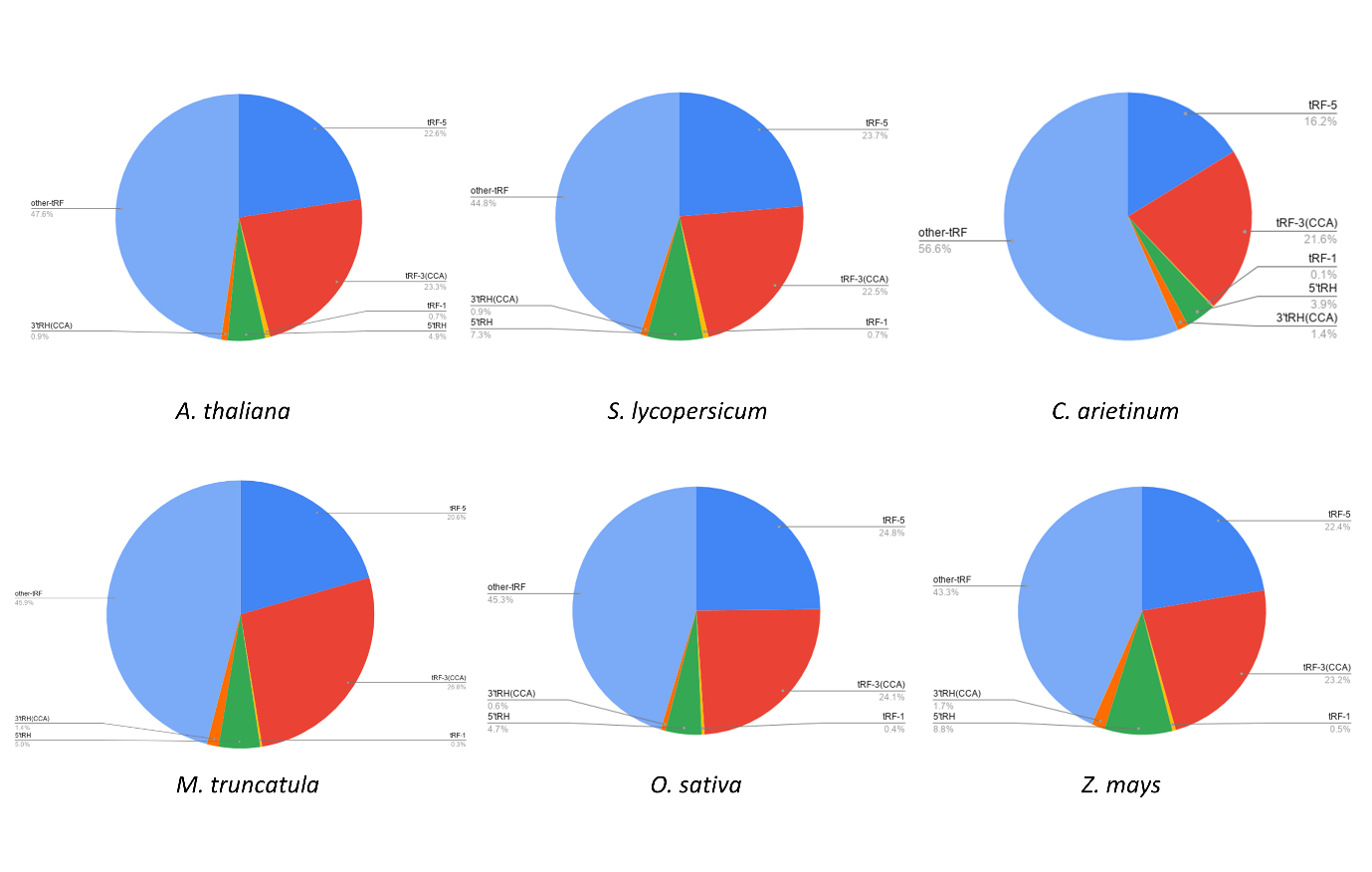

Supplementary Figure 2: The distribution of tncRNA classes in different plants.

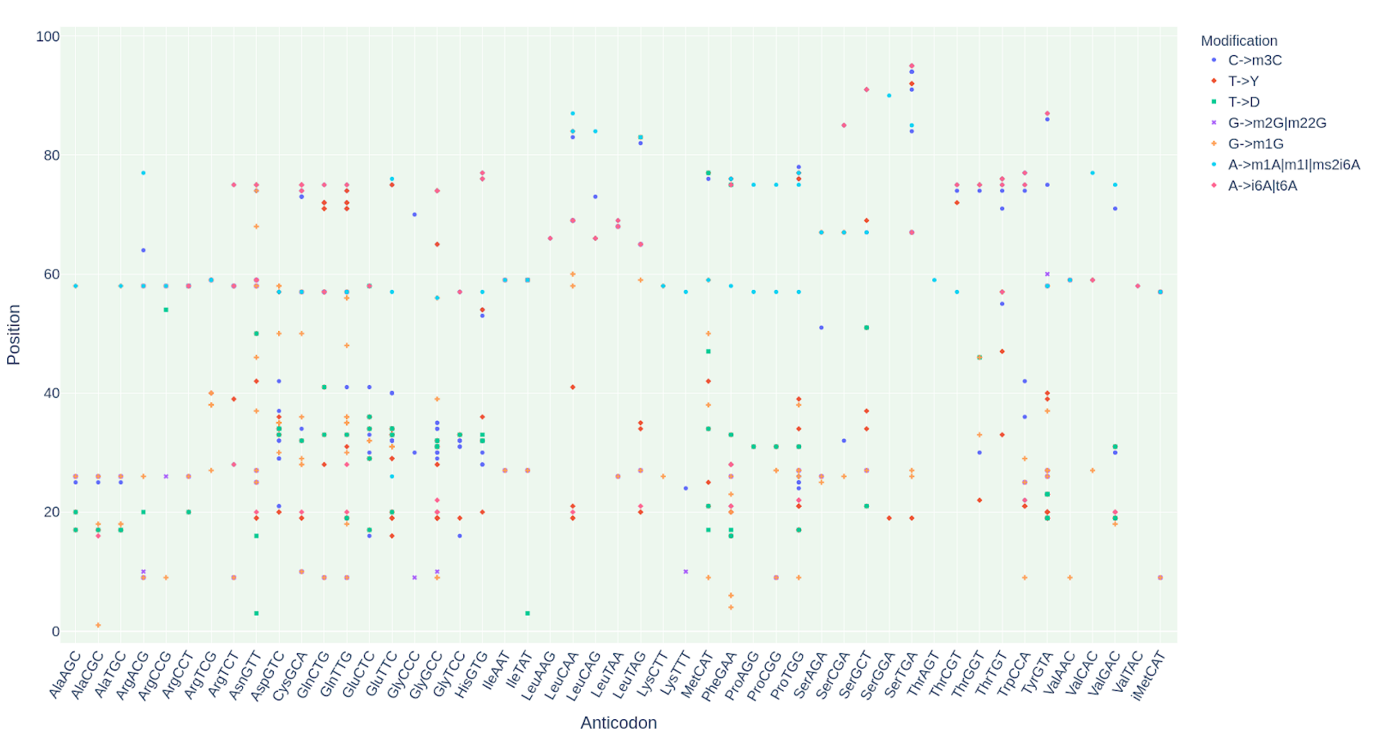

Supplementary Figure 3: The different tRNA modifications predicted by HAMR in identified tncRNAs originating from their respective tRNA anticodon in *A. thaliana*.

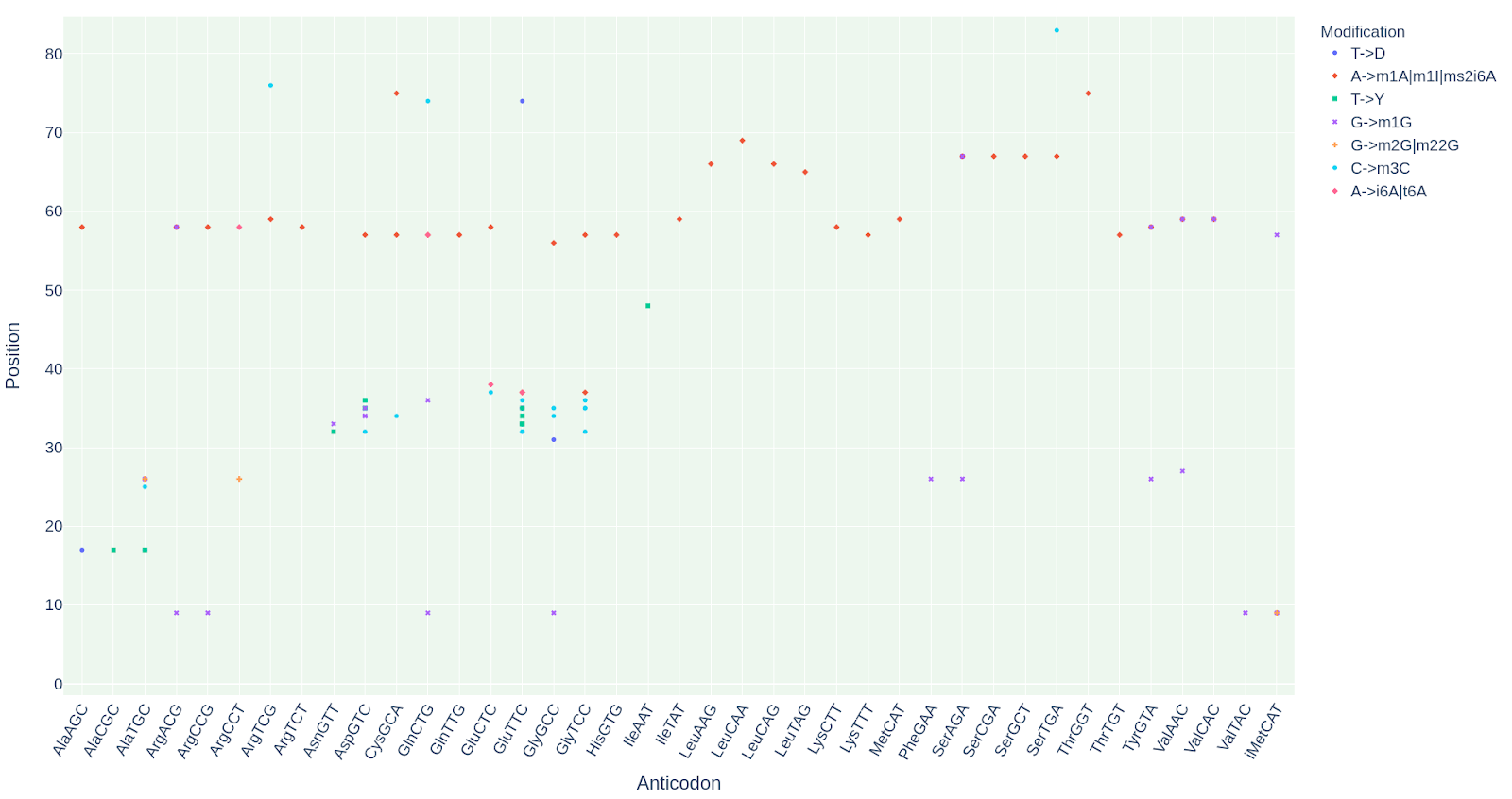

Supplementary Figure 4: The different tRNA modifications predicted by HAMR in identified tncRNAs originating from their respective tRNA anticodon in *S. lycopersicum*.

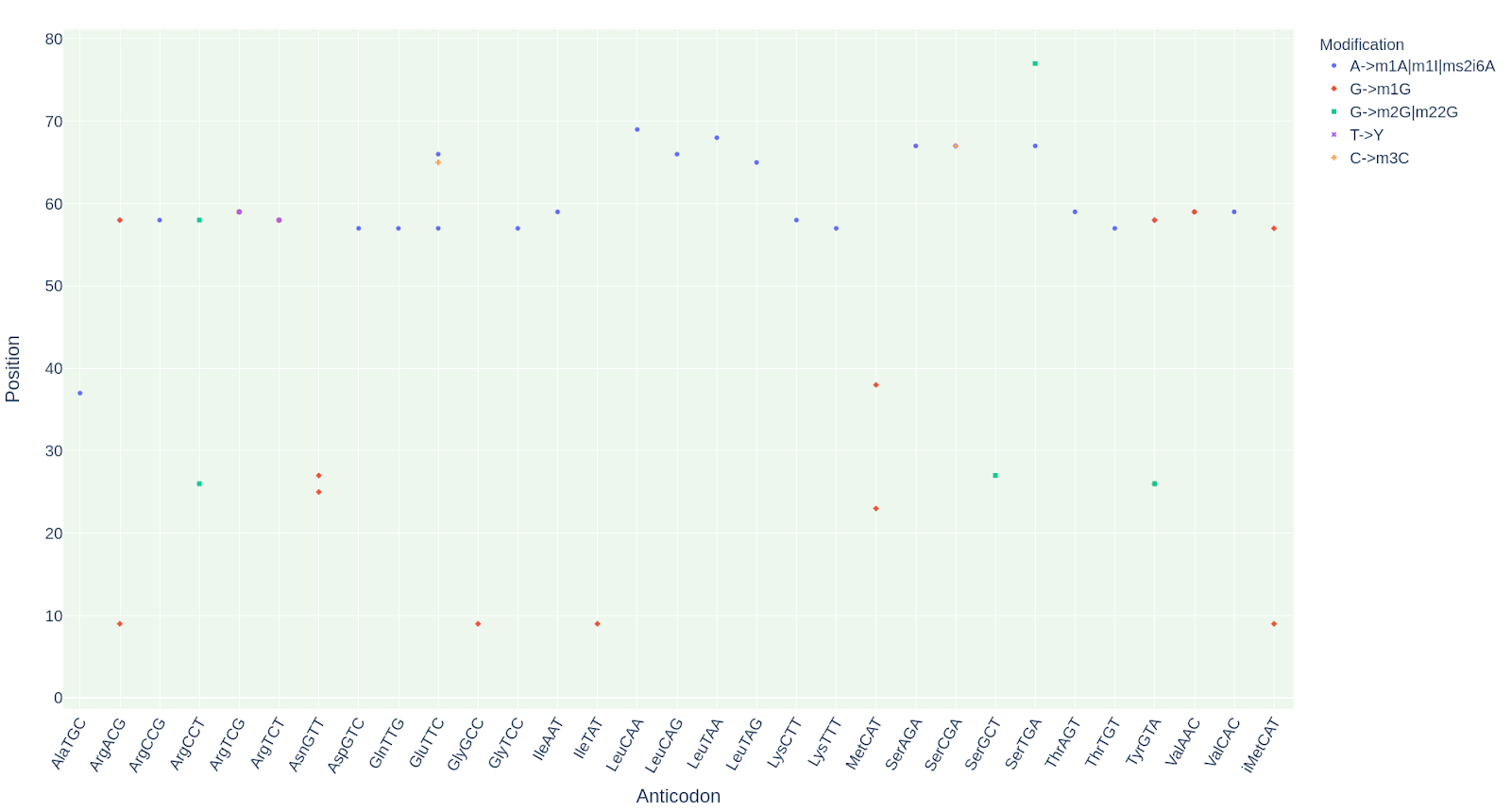

Supplementary Figure 5: The different tRNA modifications predicted by HAMR in identified tncRNAs originating from their respective tRNA anticodon in *C. arietinum*.

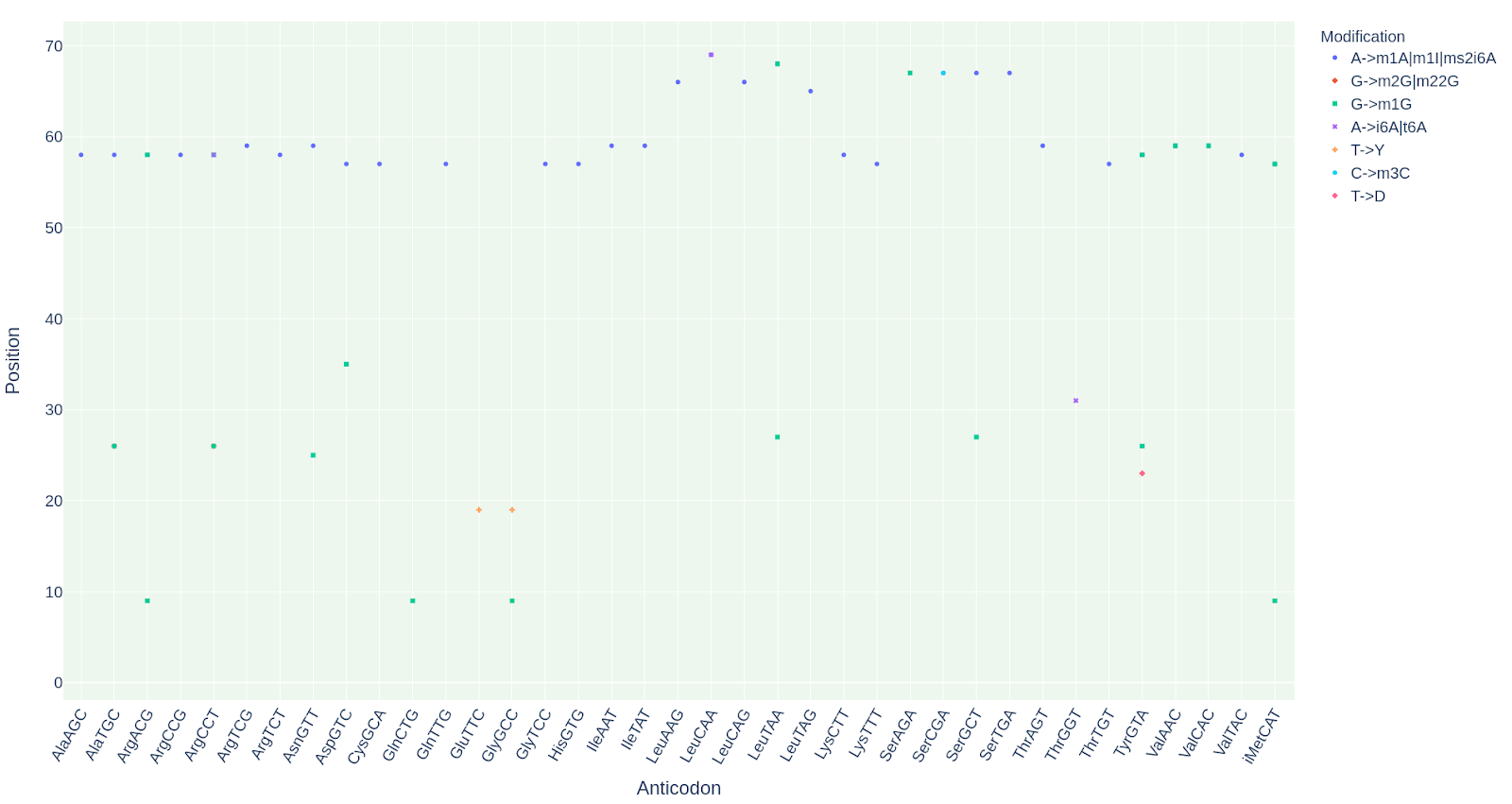

Supplementary Figure 6: The different tRNA modifications predicted by HAMR in identified tncRNAs originating from their respective tRNA anticodon in *M. truncatula*.

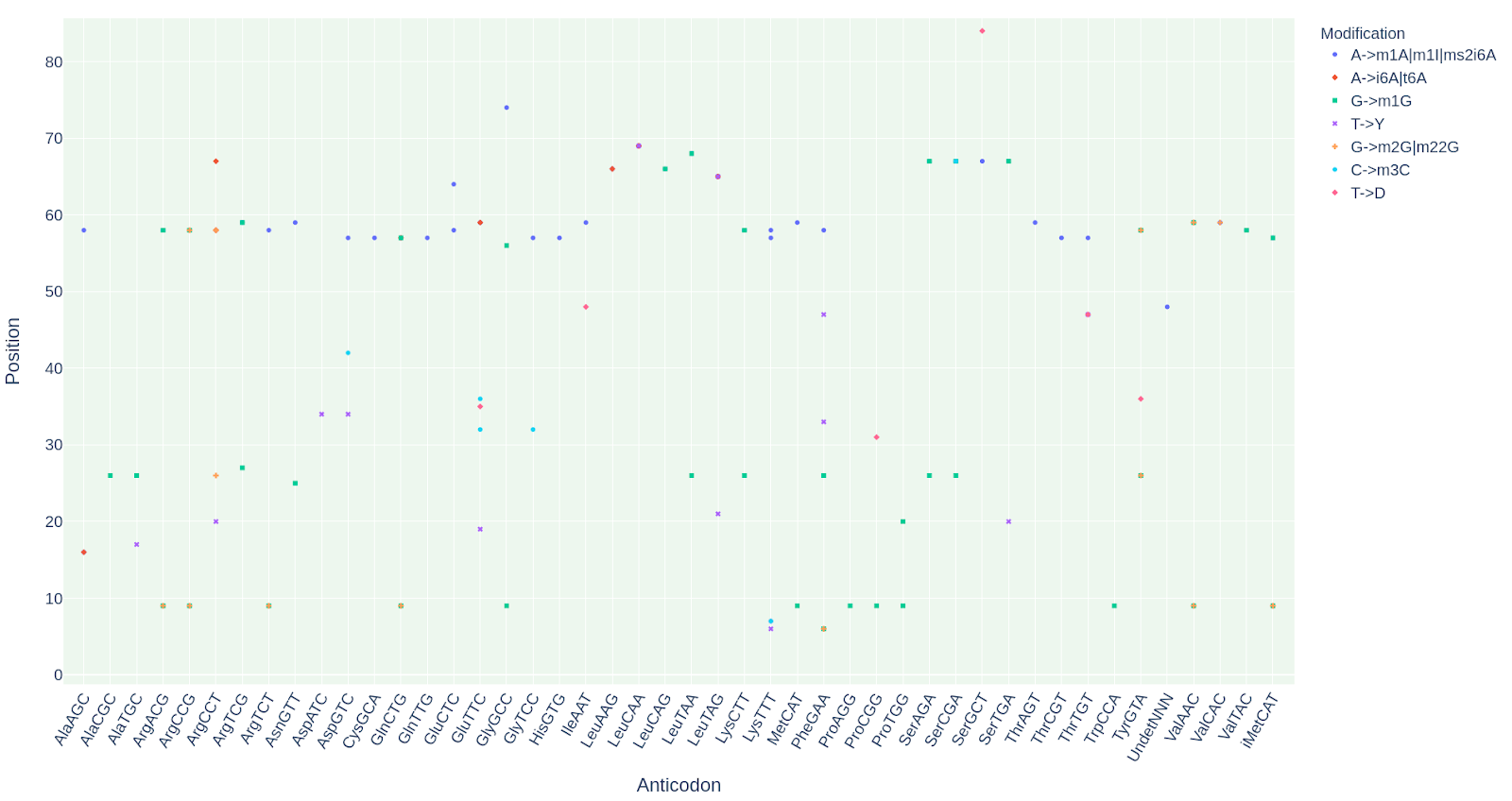

Supplementary Figure 7: The different tRNA modifications predicted by HAMR in identified tncRNAs originating from their respective tRNA anticodon in *O. sativa*.

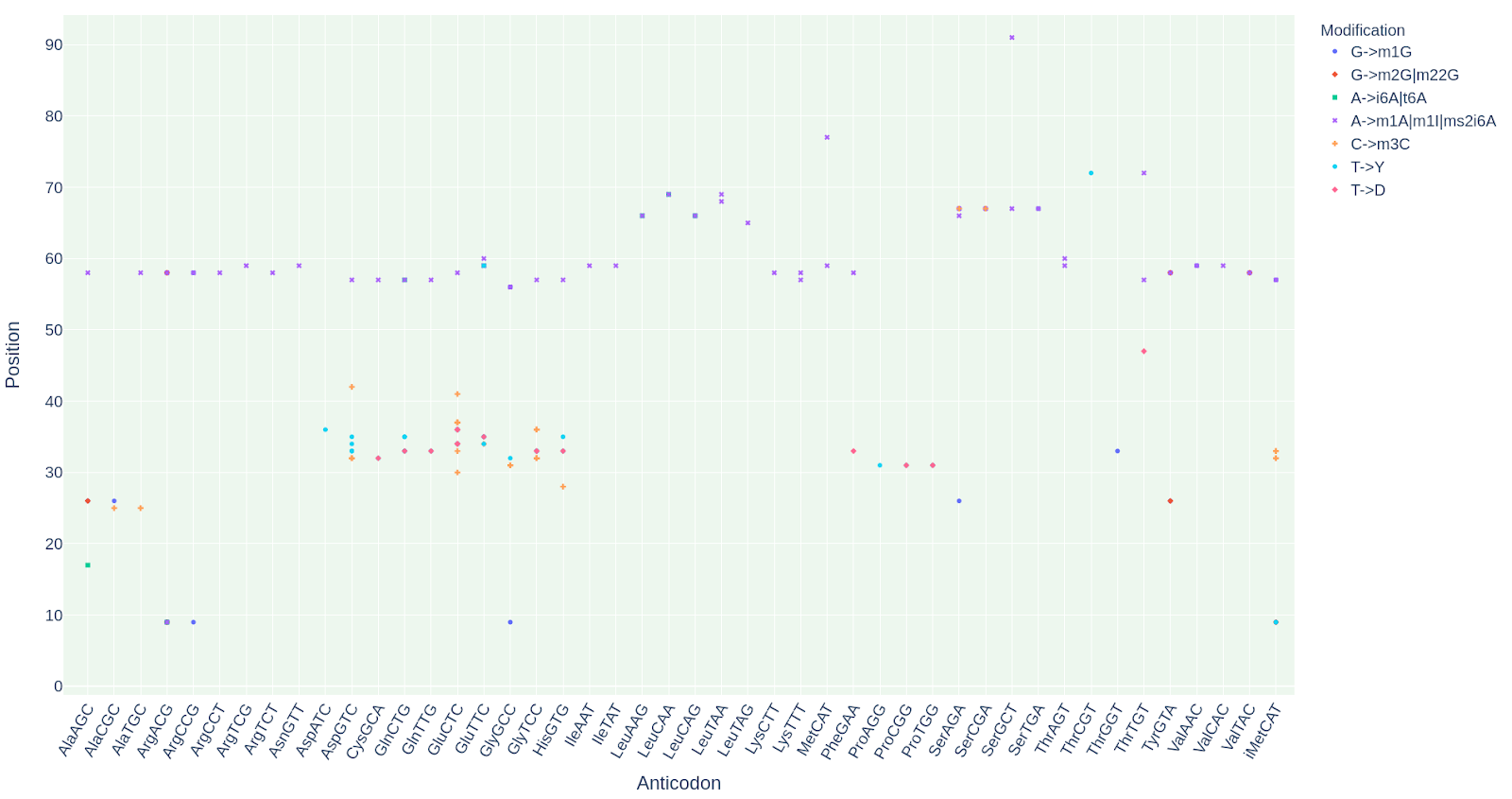

Supplementary Figure 8: The different tRNA modifications predicted by HAMR in identified tncRNAs originating from their respective tRNA anticodon in *Z. mays.*

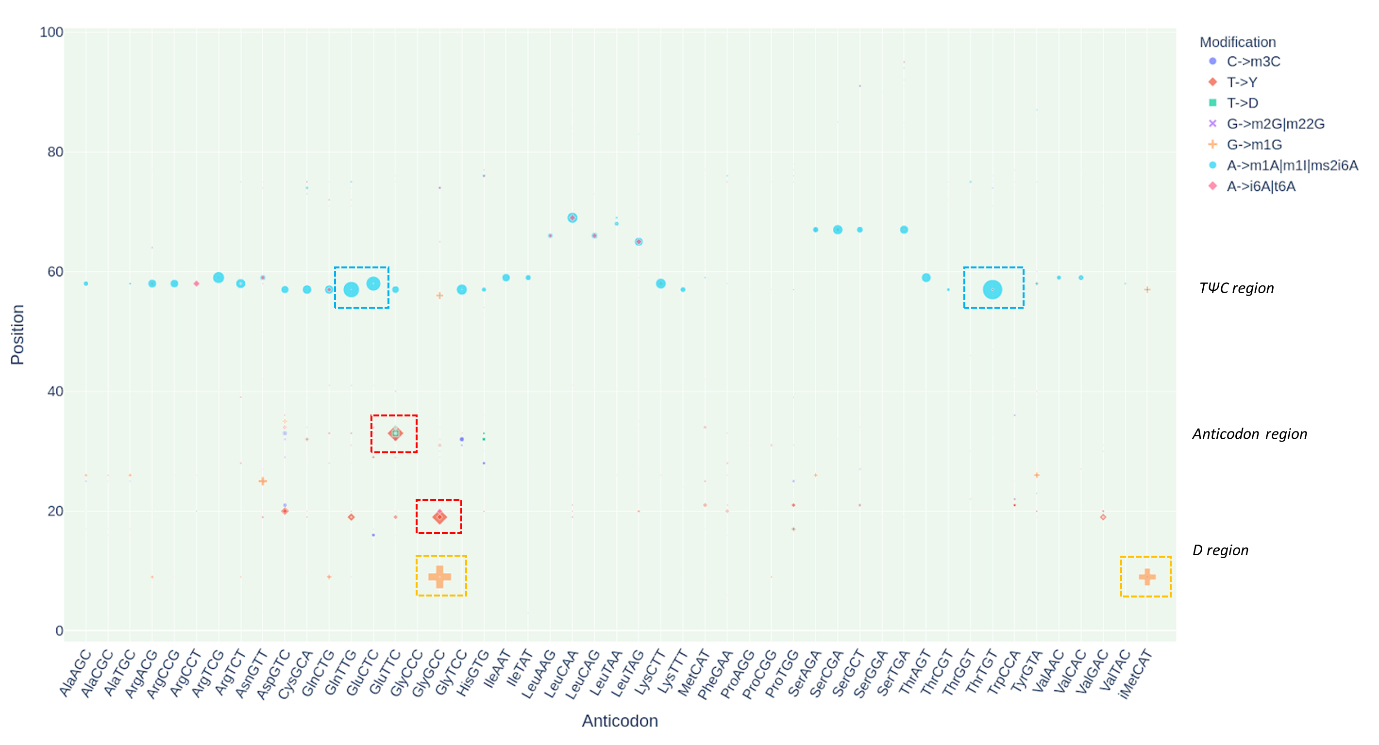

Supplementary Figure 9: The recurrent modification residues in tncRNAs originating from their respective tRNA anticodon in *A. thaliana*. The most abundant nucleoside modification has been highlight by dashed boxes.

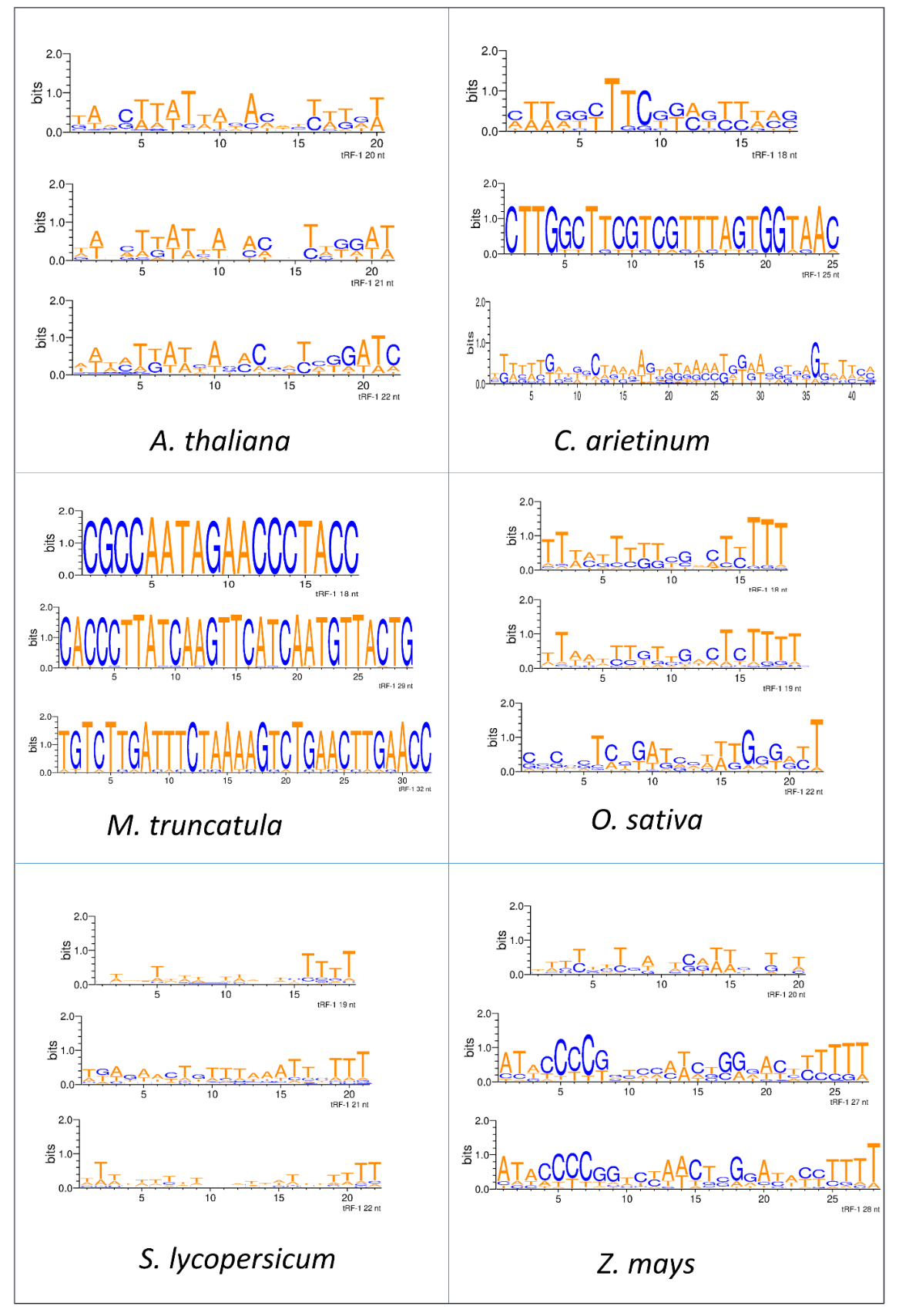

Supplementary Figure 10: Web logo representation for the top-three most recurring tRF-1 sequences (length-wise) in six plants.
